# Egg size variation in the context of polyandry: a case study using long-term field data from snowy plovers

**DOI:** 10.1101/2020.08.07.240150

**Authors:** Luke Joseph Eberhart-Hertel, Lourenço Falcão Rodrigues, Johannes Krietsch, Anne G. Eberhart-Hertel, Medardo Cruz-López, Karina Alejandra Vázquez-Rojas, Erick González-Medina, Julia Schroeder, Clemens Küpper

**Affiliations:** Max Planck Institute for Ornithology; Universidad Autónoma de Madrid; Seckenberg Biodiversity and Climate Research Centre, Ludwig-Maximilians University of Munich; Institute of Ocean Sciences and Limnology, Universidad Nacional Autónoma de México; University of Extremadura; Silwood Park Campus, Imperial College London

**Keywords:** breeding phenology, *Charadrius nivosus*, female-female competition, mating system, reproductive investment, season, senescence

## Abstract

Gamete size variation *between* the sexes is central to the concept of sex roles, however, to what extent gamete size variation *within* the sexes relates to sex role variation remains unclear. Comparative and theoretical studies suggest that, when clutch size is invariable, polyandry is linked to a reduction of egg size, while increased female-female competition for mates favors early breeding when females cannot monopolize multiple males. To understand whether and how breeding phenology, egg size, and mating behavior are related at the individual level, we studied the reproductive histories of 424 snowy plover females observed in the wild over a 15-year period. Egg size, but not polyandry, were highly repeatable for individual females. Consistent with theoretical predictions, we found that polyandrous females were the earliest breeders and that early clutches contained smaller eggs than clutches initiated later. Neither egg size nor mating behavior showed clear signs of an age-related deterioration, on the contrary, prior experience acquired either through age or local recruitment enabled females to nest early. Taken together, these results suggest that gamete size variation is not linked to mating behavior at the individual level, and, consequently, the adaptive potential of such variation appears to be limited.

**Teaser Text:** Comparative avian analyses have linked female polyandry and sex-role reversal to the production of smaller eggs: the notion being that smaller eggs allow polyandrous females to lay multiple sequential clutches quickly. Our research, however, has found that the notion is more complex at the individual level. We found that females who started breeding early in the season were more likely to be polyandrous and produce smaller eggs. Experience was also found to give older and locally-raised females a competitive advantage over younger and inexperienced females vying for early breeding opportunities. Despite this, we found limited evidence that egg size reduction gives a competitive advantage in the scramble for mates. Instead, seasonal variations in egg size can likely be attributed to other factors, such as the survival of early-hatching chicks, resource limitations during egg production, and individual differences. We found that senescence had little impact on egg size or mating behavior. Future research should also examine how seasonality, age, and mating behavior impact the reproductive fitness of males, who provide more parental care than females in many sex role reversed systems. By studying these effects in concert, we can gain a deeper understanding of sex roles in natural populations.

## INTRODUCTION

Anisogamy is a fundamental prerequisite for the concept of sex roles (e.g., Darwin 1871; Bateman 1948; Wade and Shuster 2002; Schärer et al. 2012; Lehtonen et al. 2016). In species with conventional sex roles, males, the producers of many small gametes, invest into acquiring matings to sire as many offspring as possible, whereas females, the producers of few large gametes, aim to improve the survival prospects of their offspring by obtaining critical resources and providing parental care. However, there are many species where sex roles do not conform to this conventional paradigm (reviewed in Janicke et al., 2016). Moreover, there can also be considerable gamete size variation within the sexes – however, the causes and consequences of this intrasexual variation remains unclear (Bernardo, 1996; Immler et al., 2011). For example, the largest eggs produced by some females are typically at least 50% larger than the smallest eggs produced in the same population (Christians, 2002) and occasionally the largest egg can be more than twice as heavy as the smallest viable egg (Svagelj & Quintana, 2011). Such among-individual variation in egg size may have important consequences for individual fitness and could be linked to different reproductive strategies of females. Egg size is an important life history trait under maternal control that is positively associated with offspring fitness (Smith & Fretwell, 1974; Williams, 1994; Christians, 2002; Krist, 2011). Egg size typically reflects hatchling quality: the amount of nutrients provided to the developing offspring are especially important during early life (Williams, 1994; Krist, 2011). Because resources for maternal investment during oogenesis are typically limited, among- and within-female variation in egg size may signal a trade-off between offspring quality and quantity.

In oviparous organisms, egg size represents a basic measure of maternal investment (Kaplan, 1980; Fox, 1994; Williams, 1994, 2012; Starck & Ricklefs, 1998; Moran & Emlet, 2001; Xu *et al*., 2019) and is shown to be related to inter- and intra-specific variation in several life history traits. For example, egg size is associated with developmental mode, with precocial species typically producing larger eggs than altricial species (Smith & Fretwell, 1974; Williams, 1994; Christians, 2002; Krist, 2011). Within species a substantial proportion of the egg size variation can be explained by consistent individual differences between females which are highly repeatable across breeding attempts (Christians, 2002). In some cases, intraspecific variation in egg size is related to among-female variation in reproductive strategies (Slagsvold *et al*., 1984) and is occasionally linked to discrete demographic classes such as reproductive morphs (Beck *et al*., 2022; Giraldo-Deck *et al*., 2022).

Given the strong association between egg size variation and life-history traits or maternal investment it is not surprising that there are strong temporal components related to egg size variation. For example, reproductive output is often age-dependent (Bouwhuis et al., 2009; Dingemanse et al., 2020; Hammers et al., 2012; Jankowiak et al., 2018; Lemaître et al., 2015; Reznick et al., 2004; Riecke et al., 2023; Salguero-Gómez et al., 2016; Zhang et al., 2015) whereby an individual’s performance increases over early life to a maximum, followed by senescent decline. Senescence is a consequence of age-dependent trade-offs between energy investments in reproduction at the expense of somatic repair (Kirkwood & Rose, 1991). Decreasing egg and/or clutch size can indicate reproductive senescence that reduces female reproductive performance with older ages (Reid *et al*., 2003; Beamonte-Barrientos *et al*., 2010; Dingemanse *et al*., 2020; Fay *et al*., 2021). Conversely, polyandry has been associated to female age, with older more experienced females being more likely to be polyandrous than younger females (Oring *et al*., 1994). Such observations of higher mating rates at older ages suggest that differences in individual quality or experience are related to female mating behavior (Whittingham & Dunn, 2010), which could in turn be linked to among- and within-female variation in egg size.

Another source for temporal variation in egg size is seasonality. Egg size often changes throughout the reproductive season, especially in birds (Dittmann & Hötker, 2001; Skrade & Dinsmore, 2013; Weiser *et al*., 2018; Kubelka *et al*., 2020; Verhoeven *et al*., 2020). At the population level, reproductive performance often declines with season which may coincide with declining egg sizes (Varpe, 2017; Weiser *et al*., 2018). Several processes may contribute to this trend. First, in environments with limited or seasonally dependent resources, breeding early may provide favorable conditions for raising offspring, thereby reducing the amount of parental care required and maximizing offspring survival prospects (Varpe, 2017; Weiser *et al*., 2018). Consistent with this, offspring that are born early in the season often have higher survival prospects than offspring born late in the season (Festa-Bianchet, 1988; Reznick *et al*., 2006; Plard *et al*., 2015; Cruz-López *et al*., 2017; Varpe, 2017). Second, only high quality and experienced females may be able to reproduce early in the season because of potential increased physiological stress for early egg-laying females, and/or the increased likelihood of brood failure in environments with high stochasticity, such as low food availability, inclement weather, or frequency-dependent predation risk (Borgmann *et al*., 2013; Ockendon *et al*., 2013). Therefore, less experienced or low-quality females may need to postpone their reproduction to a later date when conditions are more benign. Third, alternation between extended breeding and non-breeding periods may contribute to differences in egg and, consequently, offspring quality: when breeding is restricted to a certain period of the year, females can use the preceding non-reproductive period to amass resources for investment into egg or clutch size of their initial breeding attempt (Yohannes *et al*., 2010). As the breeding season advances, less time is available for successful breeding attempts and as a consequence females may acquire less resources for egg production resulting in fewer and or/smaller eggs in late clutches (Wallander & Andersson, 2003).

An understudied evolutionary process explaining egg size variation within species is sexual selection acting on females. Although sexual selection is predominantly considered to shape male-male competition, it is increasingly acknowledged that sexual selection also operates on females with strong female-female competition for mates (Clutton-Brock, 2007; Kraaijeveld *et al*., 2007; Fromonteil *et al*., 2023). In some species, competition over mating opportunities is stronger in females than in males, leading to a reversal of conventional sex roles (Oring & Lank, 1982; Rosenqvist, 1990; Andersson, 2004; Goymann *et al*., 2015; Fritzsche *et al*., 2021). Theoretical models have linked intrasexual competition to variation in gamete size, with more intense competition leading to smaller gametes (Wade & Shuster, 2002; Lehtonen *et al*., 2016). Notably, this relationship is not restricted to males and can also be applied to females in polyandrous species with female-female competition over mates, resulting in a reduction of egg size. For instance, at the species level, a comparative analysis of birds showed that a lineage’s egg size tends to decrease with the evolution of polyandry (Oring & Lank, 1982; Rosenqvist, 1990; Andersson, 2004; Goymann *et al*., 2015; Fritzsche *et al*., 2021) – supposedly due to the selective advantages that producing smaller eggs has on minimizing the laying period between matings (Goymann *et al*., 2015).

Andersson (2004) presented a theoretical quantitative genetics model in which intense female scramble competition (i.e., whereby females compete to find sexually receptive males to commence breeding) shapes breeding phenology and egg size variation in females. According to the model, early breeding provides substantial advantages as it allows these females enough time to pursue sequential breeding attempts or to replace a failed first attempt (i.e., "re-nesting"; Morrison *et al*., 2019). Early breeding should therefore be tightly correlated with female polyandry and, ultimately enhance the fitness of polyandrous females. Laying smaller eggs could provide another competitive edge in female-female competition over males: when clutch sizes are uniform, females that produce small eggs can complete their clutch faster than those that produce larger eggs and hence gain a competitive edge in the competition for mates especially when breeding synchrony is high and additional males are limited. A lower boundary for egg size is provided by reduced offspring viability because smaller eggs have fewer nutrients (Williams, 1994; Blomqvist *et al*., 1997; Krist, 2011; Giraldo-Deck *et al*., 2022). However, the predictions on breeding phenology, egg size, and their relationship with female mating behavior provided in Andersson’s (2004) theoretical model remain to be tested empirically at the individual level.

Among oviparous animals, shorebirds (part of the order Charadriiformes) produce some of the largest eggs in relation to body mass due to the needs of their precocial and (predominantly) nidifugous young (Lack, 1968; Rahn *et al*., 1975). Clutch size is highly uniform within shorebird species and rarely exceeds four eggs (Rahn *et al*., 1975) meaning that egg size represents an adequate proxy for maternal investment. As a clade, shorebirds also exhibit a disproportionately high prevalence of polyandry (Andersson, 2005; Colwell, 2010; Oring, 1986). Here, we investigate breeding phenology, egg size variation, and their relation with female mating strategy in snowy plovers (*Charadrius nivosus*) using a 15-year longitudinal mark-recapture dataset. The snowy plover is a long-lived shorebird (longevity record: 20 years; Colwell et al., 2017; M. A. Colwell *personal communication,* 15 March, 2021) exhibiting flexible sex-specific parental roles with sequential social monogamy and polygamy occurring in the same population (Warriner *et al*., 1986; Eberhart-Phillips *et al*., 2017). In parts of the species’ range, the breeding season can extend up to six months allowing for multiple nesting attempts following successful fledging of young or replacing failed nests due to depredation or flooding (Eberhart-Phillips, 2019; Plaschke *et al*., 2019). On average, females have higher polygamy potential and more partners per season than males, who in turn, provide more parental care than females, specifically for boods (Carmona-Isunza *et al*., 2017; Cruz-López *et al*., 2017; Eberhart-Phillips *et al*., 2017). The reason for the higher polygamy potential in females is a surplus of males in the adult breeding population (Stenzel *et al*., 2011; Carmona-Isunza *et al*., 2017; Eberhart-Phillips *et al*., 2017). Consequently, sequential polyandry, whereby females regularly desert their broods after hatching to start a new breeding attempt with another male, is commonly observed in snowy plovers (Warriner *et al*., 1986). Female desertion for re-mating declines with season in polyandrous plovers (Amat *et al*., 1999; Székely *et al*., 1999; Cruz-López *et al*., 2017; Kupán *et al*., 2021) suggesting that female-female competition for mates should be most pronounced early in the season.

To disentangle the relationships between gamete size, breeding phenology, and female mating behavior, we examined the reproductive histories of 426 snowy plover females. First, we investigated the correlates of re-nesting and female polyandry. We hypothesized that early nesting females would have higher probabilities of re-nesting or becoming sequentially polyandrous: early nesting females can rejoin the mating pool faster and therefore have higher potential to secure an additional breeding attempt compared to later nesting females. Secondly, we explored the predictors of clutch initiation date for a given female in a season. We hypothesized that early nesting females would have a competitive advantage over late nesters – either due to a direct body size advantage or an indirect advantage gained through prior experience as a previous breeder or local natal recruit. To evaluate this, we explored how body size, age, and origin (i.e., immigrant or natal recruit) were related to female nesting phenology and the likelihood of re-nesting or polyandry. Third, we examined the pattern of egg size variation in relation to season, age, and female body size. We predicted that early-laying females would produce the smallest eggs compared to later laying females due to the costs associated with intense female-female competition at the start of the season, i.e., the ‘race’ to breed early and maximize re-nesting or polyandry potential (Andersson, 2004). If reproductive senescence affects egg size variation, we predicted egg size to follow a quadratic relationship with female age. Alternatively, if female polyandry increases with age, we predicted a negative linear relationship of egg size with female age.

## MATERIALS AND METHODS

### Data collection

We studied the reproductive effort and breeding schedules of snowy plovers at Bahía de Ceuta – an important breeding site located on the coast of Sinaloa, western Mexico (23°54’N, 106°57’W). Details on the study site and population are provided elsewhere (e.g., Cruz-López et al., 2017; Eberhart-Phillips et al., 2020). In brief, we annually monitored breeding birds from mid-April until early July and collected mark-recapture data following the methods described in Székely et al. (2011). We searched for nests using telescopes and mobile hides to minimize disturbance. Upon finding a nest, we measured the length and width of each egg to the nearest tenth of a mm to determine their size (Figs. 1a, b). Using these egg dimensions, we calculated egg volume (Fig. 1c) following (Hoyt, 1979) as:

**Figure 1.**
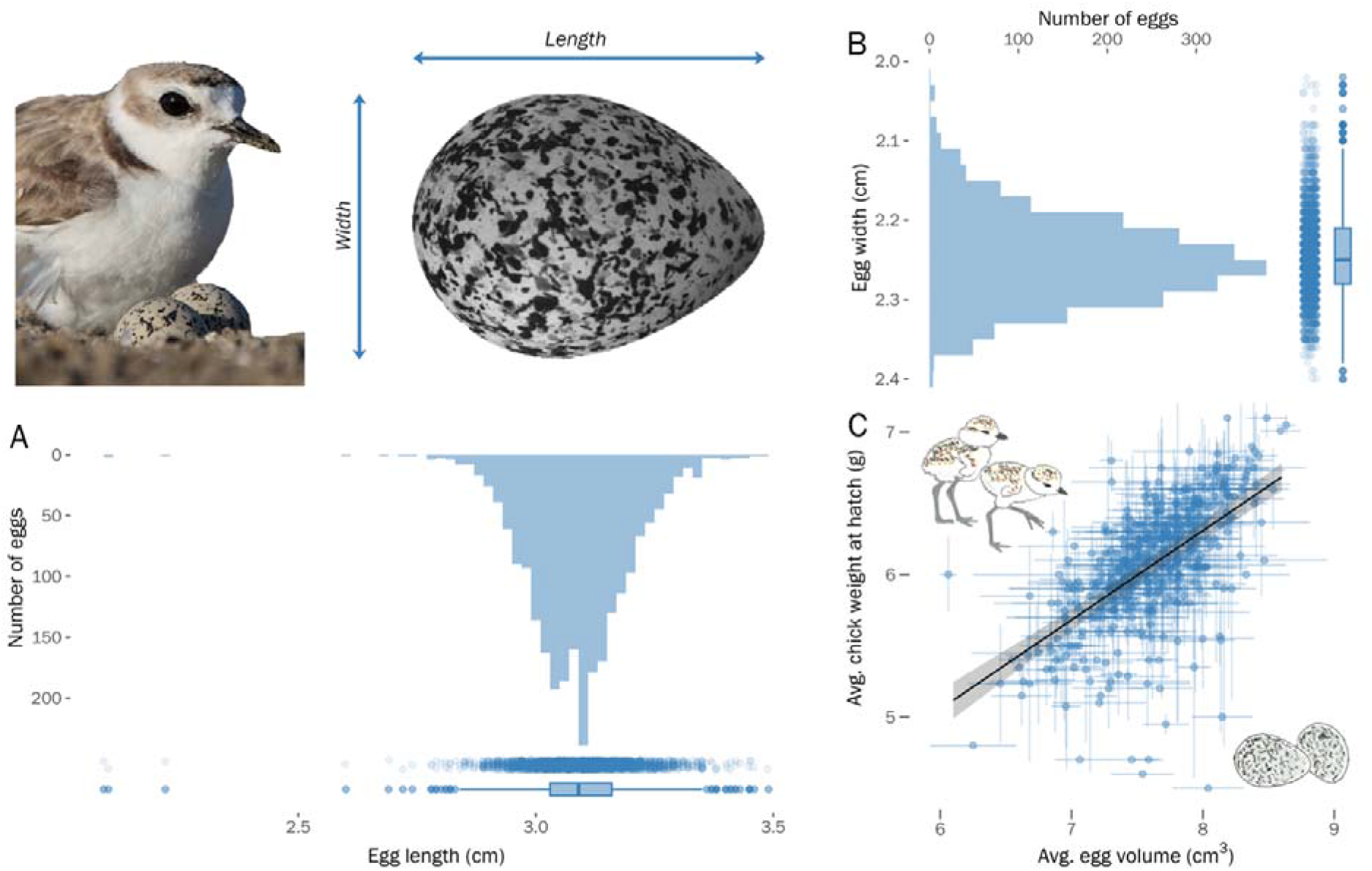
Egg size variation for snowy plovers breeding in Bahía de Ceuta, Mexico between 2006 and 2020. Egg volume was estimated using an allometric equation combining length (A) and width (B) measurements collected in the field (see *Methods*). C) Average egg volume of a clutch was highly correlated with the clutch’s average chick weight at hatch (β [95% CIs]: 0.628 [0.552–0.704]; R^2^*_marginal_* = 0.370 [0.310–0.436]). Ribbon shows the 95% CI of the model prediction, vertical and horizontal error bars visualize within clutch variation in egg volume and chick hatch weight (1 SD); see Supplementary Text D for modelling methods.

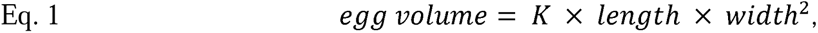

where *K* is 0.486, a volume-index constant for snowy plovers determined by (Székely *et al*., 1994) through the use of an egg volumeter (Hanson, 1954). The modal clutch size of snowy plovers is three (91% of 836 nests in our sample) and is the maximum number of eggs we have observed in this population (Eberhart-Phillips *et al*., 2020a). We regularly checked incomplete nests until the clutch was complete and assigned the age of these nests according to the date of the last egg laid (Plaschke *et al*., 2019). If the clutch was complete upon discovery and had not been incubated longer than 10 days (92.5%; *N* = 773 nests), we determined its lay date by floating the egg and estimating the stage of embryonic development. For successful clutches that had been incubated for more than 10 days (5.5%, *N* = 46 nests), we back-calculated the laying date based on the hatching date assuming an incubation period of 25 days (Plaschke *et al*., 2019). In the rare case that the nest did not hatch and we discovered it after day 10 of incubation (2%, *N* = 17 nests), we assumed that the nest was 17 days old upon discovery (i.e., the midpoint between the minimum age of 11 days and the 25-day incubation period).

We identified previously marked nesting adults based on their unique colour ring combination. We captured unmarked adults on their nests during incubation using funnel traps (Hall & Cavitt, 2012) and assigned a unique colour ring combination for subsequent recognition. In the rare circumstance when we were unable to identify parents before hatching, we attempted to capture parents while they tended the chicks. As snowy plovers only show a small degree of sexual dimorphism, we determined the sex of all captured plovers in the field through a combination of plumage characteristics (Argüelles-Tico *et al*., 2015), time of capture (i.e. females typically incubate during the day and males during the night; <u>Vincze *et al*., 2017</u>), and other behavioral cues (e.g., sex-specific brood care; Kupán et al., 2021). For a subset of adults (57.5%), we confirmed sex molecularly from DNA extracted from blood samples by PCR amplification of Z and W specific DNA regions with two sex-typing markers: P2/P8 and Calex-31 (Griffiths *et al*., 1998; Küpper *et al*., 2009b; dos Remedios *et al*., 2010). Over our 15-year study we conducted 1168 captures of individuals that had been molecularly sex-typed: in only 2.4% of these captures did we mis-identify the sex of an individual using field-based cues.

We visited known active nests every four or five days to determine the status of the nest (e.g., active, depredated, etc.) until the 20^th^ day after egg laying and thereafter daily until the eggs hatched or failed. We weighed the chicks shortly after hatching (879 [84.7%] within 24 hours after hatching, 159 [15.3%] during the second day after hatching) and marked them with an alphanumeric metal and a single color ring for subsequent identification in the chance that these individuals would be recruited into the breeding population as adults in future years.

For the years 2006 to 2016 all longitudinal data collected has been compiled as part of the *CeutaOPEN* project – an open-access database for individual-based field studies in evolutionary ecology and conservation biology (Eberhart-Phillips *et al*., 2020a). We accessed these data directly from the open source repository (Eberhart-Phillips *et al*., 2020b) and supplemented them with data from four additional field seasons: 2017–2020. Please refer to our RMarkdown vignette that connects to *CeutaOPEN* and reproduces all analytical methods and results presented below (Supplementary File 1). All of our field activities were performed and permitted in accordance with the approved ethical guidelines outlined by SEMARNAT (Secretariat of Environment and Natural Resources, Mexico).

### Statistical Analyses

To evaluate the relationship between female breeding behavior, nesting phenology, and egg size, our investigation implemented three step-wise analyses that: 1) established the factors related to a female’s potential to have multiple breeding attempts either through polyandry or re-nesting, 2) identified the key female traits linked to lay date, and 3) dissected the elements explaining within- and among-female variation in egg volume investment. A central trait of interest in these three analyses was female age, as we needed to acknowledge age-dependent variation in the repeated measures of individuals over the 15-year period of data collection. We first present our statistical methods for estimating the age of females that entered the population as adults, then we describe our models for investigating the interplay between female breeding behavior, nesting phenology, and egg size.

#### Statistical estimation of age for females first encountered as adults

To estimate the ages of unknown individuals in our marked population, we conducted a capture-mark-recapture analysis using the ‘Bayesian Survival Trajectory Analysis’ (BaSTA) package in R (v1.9.4, Colchero et al., 2012), which uses a Bayesian hierarchical framework to fit parametric survival functions of the marked population while accounting for imperfect detection (see Supplementary Text A for details). In short, the capture histories of unknown-age individuals (90%, *N =* 405) are cast against the survival function of the known-age population (10%, *N* = 45) to produce posterior age estimates. To acknowledge uncertainty in the BaSTA age estimate of an unknown age individual: we ran bootstrap simulations for each of the following models 1000 times, with every iteration randomly drawing a birth year estimate for unknown aged individuals from their posterior distributions provided by BaSTA. For each simulation, we evaluated the influence of birth year uncertainty by examining the effect size distribution of the 1000 bootstraps in relation to the 95% confidence interval for effect sizes of the original model that used the median birth year estimate from BaSTA. Furthermore, to test the accuracy of the BaSTA age estimates, we implemented a cross-validation simulation which iteratively estimated the age of each known-age female under an *in silicio* scenario whereby their age was unknown. The simulation confirmed that the BaSTA age-estimates were adequately accurate, with 51% of estimated ages being correct and a combined 82% of estimated ages being within 1-year of the true age (Fig. S1) – a notable result considering that our mark-recapture sample is from a wild and highly vagile avian species. Moreover, BaSTA age-estimates that deviated from reality were slightly biased towards an under-estimation (i.e., estimated to be younger; Fig. S1), hence making our age-specific models more conservative. Lastly, we considered a time-since marking capture-mark-recapture model and determined that the recapture probability of our dataset was not biased by individual heterogeneity in post-marking processes (see Supplementary Text B for details).

A key methodological issue for studying age related processes in wild populations is that stochastic extrinsic mortality reduces the frequency of individuals in older age classes, hence making it challenging to disentangle between- vs. within-individual age-dependent variation – a phenomenon known as “selective disappearance” (van de Pol & Verhulst, 2006a; Nussey *et al*., 2008). Investigations using longitudinal data to test for age-related processes are particularly suitable for this task, as they can control for the confounding effects of selective disappearance through repeated measures of individuals as they age (van de Pol & Verhulst, 2006a; Nussey *et al*., 2008; Dingemanse *et al*., 2020). We controlled for selective appearance and disappearance of females differing in the trait of interest by fitting ‘first observed age’ and ‘last observed age’ as fixed effects – a method that estimates between-individual age effects introduced by selective disappearance and appearance (Dingemanse et al., 2020; van de Pol & Verhulst, 2006a, 2006b). We also modelled within-individual age effects by fitting linear and quadratic forms of a within-group deviation score for age (henceforth ‘age-deviance’), calculated for individual *i* at age *j* as: *age_ij_* – [*first observed age*]*_i_* (van de Pol & Verhulst, 2006a; Snijders & Bosker, 2011) – essentially describing the number of years since the bird’s arrival to the population. To ensure that intercepts of our age-dependent models represented the reproductive performance for the earliest age at reproduction (i.e., age 1 in snowy plovers; Page et al., 2009), we fitted age as ‘*age* – 1’ – otherwise it would represent reproduction at age 0, which is an empirically meaningless estimate. Moreover, we followed common statistical approaches to investigate age-related processes in birds (e.g., Bouwhuis et al., 2009, 2010; Dingemanse et al., 2020; Graham et al., 2019; Herborn et al., 2016; Schroeder et al., 2012) by fitting a quadratic function of age to model age-specific trends. Lastly, in all models, tarsus length was included as a fixed effect proxy of female structural size, and was averaged over an individual’s lifetime of repeated measurements, i.e., our *a priori* expectation was that tarsus length is static throughout adult life and that any variation in this trait was due to measurement error.

#### Modelling seasonal variation in polyandry potential (“Polyandry model”)

Our sample for studying seasonal polyandry dynamics included 424 females for which the identity of their mates had been verified through observation. We defined observed polyandry as a binomial variable that scored an individual as being monogamous or polyandrous each year based on our observations of them having one or multiple breeding partners, respectively (see Fig. S2 for an example of the sampling distribution). Overall, we observed 90 cases of polyandry from 74 females over the 15-year period (annual average incidence of observed polyandry: 13.5%, range: 0–25.6%). To assess the relationship between the likelihood of polyandry and lay date, age, female structural size, and maternal investment, we fitted a binomial linear mixed effects model that tested the likelihood of polyandry predicted by the fixed effects of first lay date, age-deviance (see above), first observed age, tarsus length, and the average egg-volume (of the first clutch of the season). We included individual ID and year as random effects.

#### Modelling seasonal variation in re-nesting potential (“Re-nesting model”)

Our sample for studying seasonal re-nesting dynamics included 177 females for which the fate of their first nest of the season had been verified as a failure. We defined re-nesting as a binomial variable that scored an individual as being a re-nester or a single-nester each year based on our observations of them attempting to re-nest after the loss of their first clutch or not, respectively. Overall, we observed 62 cases of re-nesting from 53 females over the 15-year period following a failed attempt (annual average incidence of observed re-nesting: 28.6%, range: 0–58.3%). To evaluate re-nesting dynamics, we fitted a binomial linear mixed effects model that tested the likelihood of re-nesting predicted by the fixed effects of first lay date, age-deviance, first observed age, and tarsus length. We included individual ID and year as random effects.

#### Modelling individual variation in lay date (“Lay date model”)

We modeled the effect of age on a female’s first lay date using a univariate mixed-effect structure that included age-deviance, age-deviance-squared, first observed age, and last observed age as fixed covariates, and individual and year as random intercepts. Furthermore, we included average tarsus length as a fixed covariate in addition to recruitment status as a two-level fixed-effect that described whether a breeding female hatched locally (“local recruit”) or was first encountered as an adult of unknown origin (“immigrant”). Our sample for studying lay date dynamics used the same nest-level sample as the polyandry model above, however, as we were interested in how the recruitment status of an individual influenced breeding phenology, we excluded data from 2006 as this was the first year when we began individual marking of all birds in the population. This subset resulted in 565 nests from 375 females. We visualized the distribution of lay dates to confirm normality and to assess the population-level variance in breeding schedule – an indication of inter-female breeding asynchrony and the intensity of female-female competition for mates (Andersson, 2004).

#### Modelling individual variation in egg volume (“Egg volume model”)

Our sample for studying egg volume dynamics included 2393 eggs from 836 nests belonging to 424 females. 56 (13.2%) females had three or more years of repeated measures (Fig. S2), 83 (19.6%) had two years of repeated measures, and 285 (67.2%) were measured in a single year. Furthermore, 43 (10.1%) individuals in our sample were marked as hatchlings but later recruited as breeding adults in subsequent years (“known age”), with the remaining 381 (89.9%) individuals being initially marked as adults (“unknown age”). We modelled within-individual age effects on egg volume by fitting a univariate mixed-effect model that included linear and quadratic forms of local tenure, and assessed between-individual age effects by additionally incorporating first observed age and last observed age as fixed covariates. Female tarsus length was also included as a covariate to control for female body size, and we incorporated a quadratic function of lay date to assess seasonal variation in egg volume as several shorebird studies report seasonal increases (Skrade & Dinsmore, 2013; Kwon *et al*., 2018) or decreases in egg volume (Dittmann & Hötker, 2001; Skrade & Dinsmore, 2013; Kwon *et al*., 2018; Kubelka *et al*., 2020; Verhoeven *et al*., 2020). We also included clutch size in our model to control for this alternate source of variation in maternal investment. Because clutch size variation in snowy plovers is constrained to 1, 2, or 3 eggs and 1- or 2- egg nests are rare in our sample (collectively 9% of nests, *N =* 75), we modelled clutch size as a binary trait (i.e., “3-eggs” vs. “1- or 2-eggs”). To disentangle within- from between-individual effects of lay date on egg size, we used the same logic as with age above: first lay dates of all individuals each year represented the between-individual seasonal effect, whereas the deviation in first lay dates of an individual relative across all successive breeding attempts for a given season represented the within-individual seasonal effect. We included random intercepts for nest, individual ID, and year, and assumed a Gaussian error distribution for egg volume.

*Evaluating effect sizes and uncertainty*—We used packages “lme4” (Bates *et al*., 2015), “rptR” (Stoffel *et al*., 2017) and “partR2” (Stoffel *et al*., 2020) in R version 4.3.1 (R Core Team, 2023) to conduct our statistical modelling and assessed homoscedasticity by visually examining residuals (see Fig. S2). For each of the four mixed-effect models described above, we evaluated the uncertainty in our parameter estimates by simulating 1000 parametric bootstraps via the “partR2::partR2” function (Stoffel *et al*., 2020). Likewise, we derived nest-, individual-, and year-level repeatabilities (i.e., intra-class correlations) by simulating 1000 parametric bootstraps of the four mixed-effect models using “rptR::rpt”. We report fixed effects as standardized regression coefficients (i.e., beta weights) and repeatability as the ‘adjusted repeatability’ – interpreted as the repeatability of a given hierarchical group after controlling for fixed effects (Nakagawa & Schielzeth, 2010).

## RESULTS

### Sample summary

The modal clutch size of the 836 clutches was 3 eggs (761 nests, 91%; 2-eggs: 72 nests, 8.6%, 1-egg: 3 nests, 0.4%). Average egg length was 3.09 cm (0.11 cm SD, range: 2.09–3.49 cm, Fig. 1a) and width was 2.24 cm (0.05 cm SD, range: 2.02–2.40 cm, Fig. 1b), which translated into an average egg volume of 7.58 cm^3^ (0.48 cm^3^ SD, range: 5.10–8.99 cm^3^). The average egg volume of a clutch strongly predicted the average hatch weight of the subsequent brood (Figs. 1c, Table S1, see Supplementary Text C for methods). Based on BaSTA’s estimated birth year, 184 of the 381 unknown-age females in our sample were first observed nesting at age one (48.3%), 120 at age two (31.5%), 71 at age three (18.6%), five at age 4 (1.3%), and one at age 5 (0.3%). Of the 43 locally hatched females in our sample, 29 first nested at age one (67.4%), six were first observed nesting at age two (14.0%), two at age 3 (4.7%), three at age 4 (7.0%), three at ages 5, 7, and 8, respectively (7.0%). Female tarsus size was not related to their annual apparent survival (see Supplementary Text D for details). The average local tenure of all females in the sample was 1.57 years (2.15 SD) with an average age span of 3.12 years (2.03 SD, median: 3, range: 1–14 years) and an average of 1.56 years of observed ages per female (1.04 SD, median: 1, range: 1–8 age-specific observations). Females in our sample were typically observed nesting every consecutive year since their first observation, however, some individuals skipped years or breed elsewhere (Fig. S2; average yearly interval between nesting attempts = 1.07, 0.27 SD). On average, females made 1.43 (0.56 SD) nesting attempts per season (median = 1, range 1 to 3). The number of hatched chicks produced by females in years when they were polyandrous, was 2.19 (1.83–2.55 95% CI) chicks more than in years when they were monogamous (Fig. S5; *N* = 14 years, 424 females, 836 nests).

### Seasonal variation in polyandry and re-nesting potential

A female’s probability of being polyandrous in a given season was strongly dependent on the date she laid her first nest, with early breeders being more likely to be polyandrous than late breeders (Figs. 2a, Table 1). Likewise, a female’s likelihood of re-nesting following a failed attempt decreased with advancing first lay date (Table 2). We found no effect of female tarsus length on the likelihood of being polyandrous (Table 1) or re-nesting (Table 2). We found a potential effect of age at first breeding on the likelihood of re-nesting, with females that initiated local breeding at an older age being the most likely to re-nest (Table 2). This effect was not found in the polyandry model (Table 1). We found no evidence that polyandry or re-nesting potential was related to age since first breeding (i.e., age-deviance; Table 1). The lay date distribution of polyandrous females was bimodal, with peaks in the first and second nests occurring 11.1 days before and 29.4 days after the unimodal seasonal peak for monogamous females (Fig. 2b). Similarly, the lay date distribution of re-nesting females was also bimodal, with peaks in the first and replacement clutches occurring 9.8 days before and 28.6 days after the unimodal seasonal peak for single nesters. There was no association between egg size and the likelihood of being polyandrous (Table 1). Females had low repeatability among years in polyandry (Table 1) and re-nesting (Table 2). When pooled together to assess multi-clutching behavior (i.e., the probability of simply laying two nests or one regardless of the outcome of the first nest or the identity of the second male), females had higher repeatability than when polyandry or re-nesting were assessed separately (Table S3).

**Figure 2.**
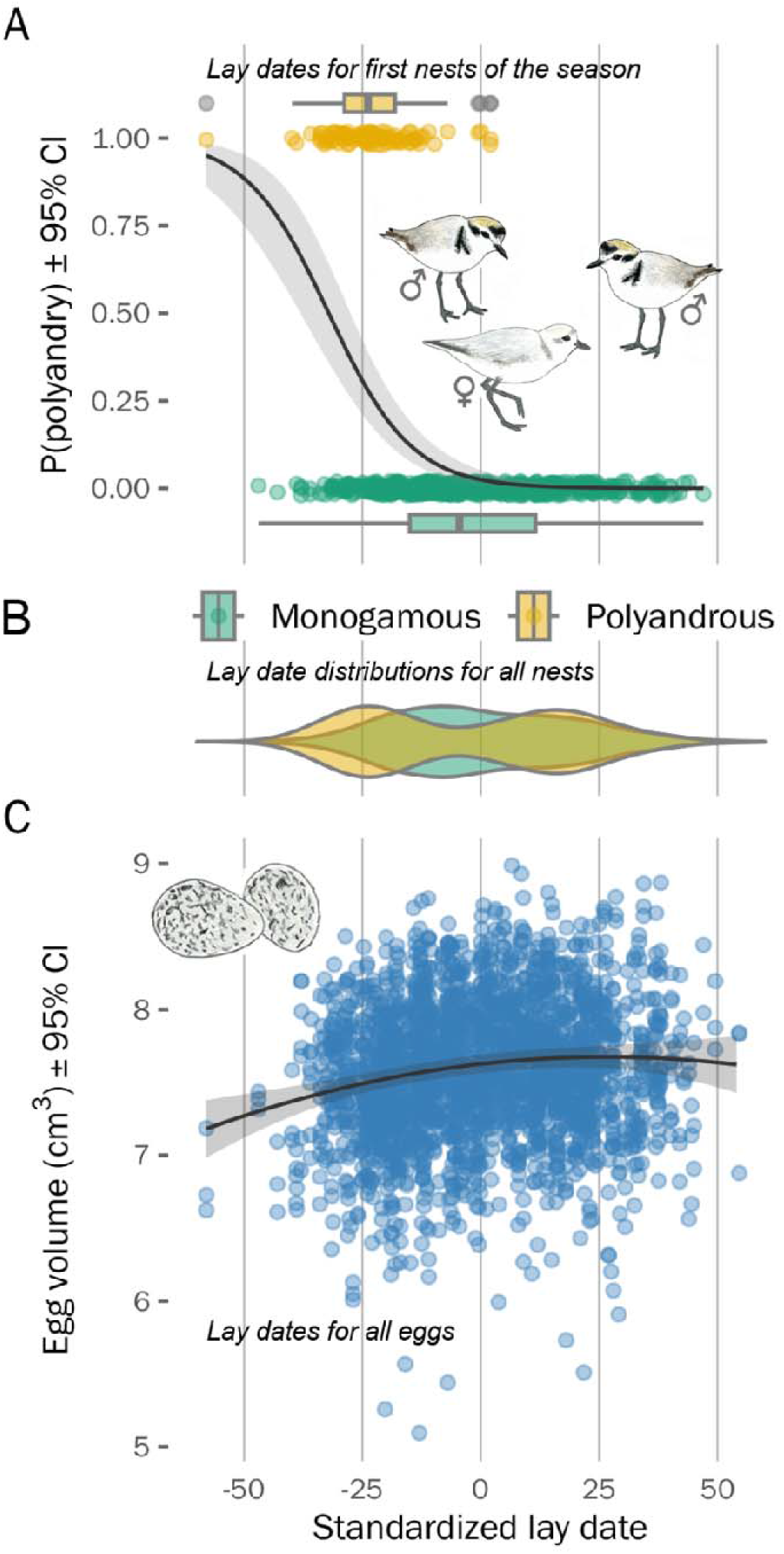
Phenology of mating behaviors, egg laying, and egg size in 424 female snowy plovers breeding at Bahía de Ceuta. A) Relationship between polyandry potential and lay date of a female’s first nest of the season. Each datum is the lay date of an individual’s first nest and their observed local mating behavior of each year. B) Lay date distributions of all nests for females that were polyandrous (yellow; 90 nests from 74 females) or monogamous (green; 571 nests from 402 females). C) Seasonal variation in egg volume – trend shows the between-individual polynomial function of the model prediction. Each datum is an egg’s volume (cm^3^) and lay date. Late date is standardized for each year across all panels.

**Table 1.** Predictors of a female’s polyandry probability in a given year: the lay date and average egg volume of their first nest of the year, their age-deviance (i.e., years since first local breeding attempt), their age at first breeding, and their structural body size. Fixed effect sizes shown are standardized estimates on the logit scale.

| *Polyandry probability* | *Mean estimate* | *95% confidence interval* |
| --- | --- | --- |
| Fixed effects $\beta$ (logit standardized) | | |
| First nest lay date | -2.12 | [-2.87, -1.68] |
| Avg. egg volume of first nest | 0.00 | [-0.29, 0.29] |
| Age-deviance | 0.00 | [-0.28, 0.26] |
| First breeding age | 0.22 | [-0.05, 0.5] |
| Female tarsus length | 0.19 | [-0.09, 0.5] |
| Partitioned $R^2$ | | |
| Total Marginal $R^2$ | 0.34 | [0.24, 0.46] |
| Total Conditional $R^2$ | 0.36 | [0.25, 0.54] |
| First nest lay date | 0.33 | [0.22, 0.45] |
| Avg. egg volume of first nest | 0.00 | [0, 0.2] |
| Age-deviance | 0.00 | [0, 0.2] |
| First breeding age | 0.01 | [0, 0.2] |
| Female tarsus length | 0.00 | [0, 0.2] |
| Random effects $\sigma^2$ | | |
| Individual | 0.07 | [0, 1.78] |
| Year | 0.16 | [0, 0.57] |
| Adjusted repeatability $r$ | | |
| Individual | 0.01 | [0, 0.17] |
| Year | 0.02 | [0, 0.06] |
| Sample sizes $n$ | | |
| Years | 14 |  |
| Individuals | 424 |  |
| Observations (i.e., Nests) | 661 |  |

**Table 2.** Predictors of a female’s re-nesting probability in a given year: the lay date of their first nest of the year, their age-deviance (i.e., years since first local breeding attempt), their age at first breeding, and their structural body size. Fixed effect sizes shown are standardized estimates on the logit scale.

| *Re-nesting probability* | *Mean estimate* | *95% confidence interval* |
| --- | --- | --- |
| Fixed effects $\beta$ (logit standardized) | | |
| First nest lay date | -1.79 | [-5.79, -1.31] |
| Age-deviance | 0.20 | [-0.19, 0.73] |
| First breeding age | 0.41 | [0.06, 1.7] |
| Female tarsus length | -0.09 | [-0.47, 0.29] |
| Partitioned $R^2$ | | |
| Total Marginal $R^2$ | 0.42 | [0.29, 0.61] |
| Total Conditional $R^2$ | 0.44 | [0.3, 0.96] |
| First nest lay date | 0.40 | [0.27, 0.6] |
| Age-deviance | 0.01 | [0, 0.43] |
| First breeding age | 0.03 | [0, 0.44] |
| Female tarsus length | 0.00 | [0, 0.43] |
| Random effects $\sigma^2$ | | |
| Individual | 0.39 | [0, 33.96] |
| Year | 0.00 | [0, 1.67] |
| Adjusted repeatability $r$ | | |
| Individual | 0.03 | [0, 14.04] |
| Year | 0.00 | [0, 0.21] |
| Sample sizes $n$ | | |
| Years | 13 |  |
| Individuals | 177 |  |
| Observations (i.e., Nests) | 209 |  |

### Individual variation in lay date

Females showed moderate repeatability in their first lay date among years (Table 3). We found strong support for the effect of origin on first lay date: females that locally hatched and later recruited into the breeding population initiated nests 7.80 days earlier (95% CI: [5.09, 10.50]) on average compared to conspecifics whose origin was unknown (Table 3 and Fig. 3b). The next strongest fixed effect was within-individual age: young females laid nests later compared to older females with lay date advancing by ∼2.17 days per year until age six (95% CI: [1.41, 2.93]; Fig. 3a), after which the uncertainty in the trend became unwieldly in the oldest age classes of our sample (Fig. 3a). Notably, female size did not affect lay date (Table 3 and S3a), nor did between-individual effects of first or last age at breeding (Table 3).

**Figure 3.**
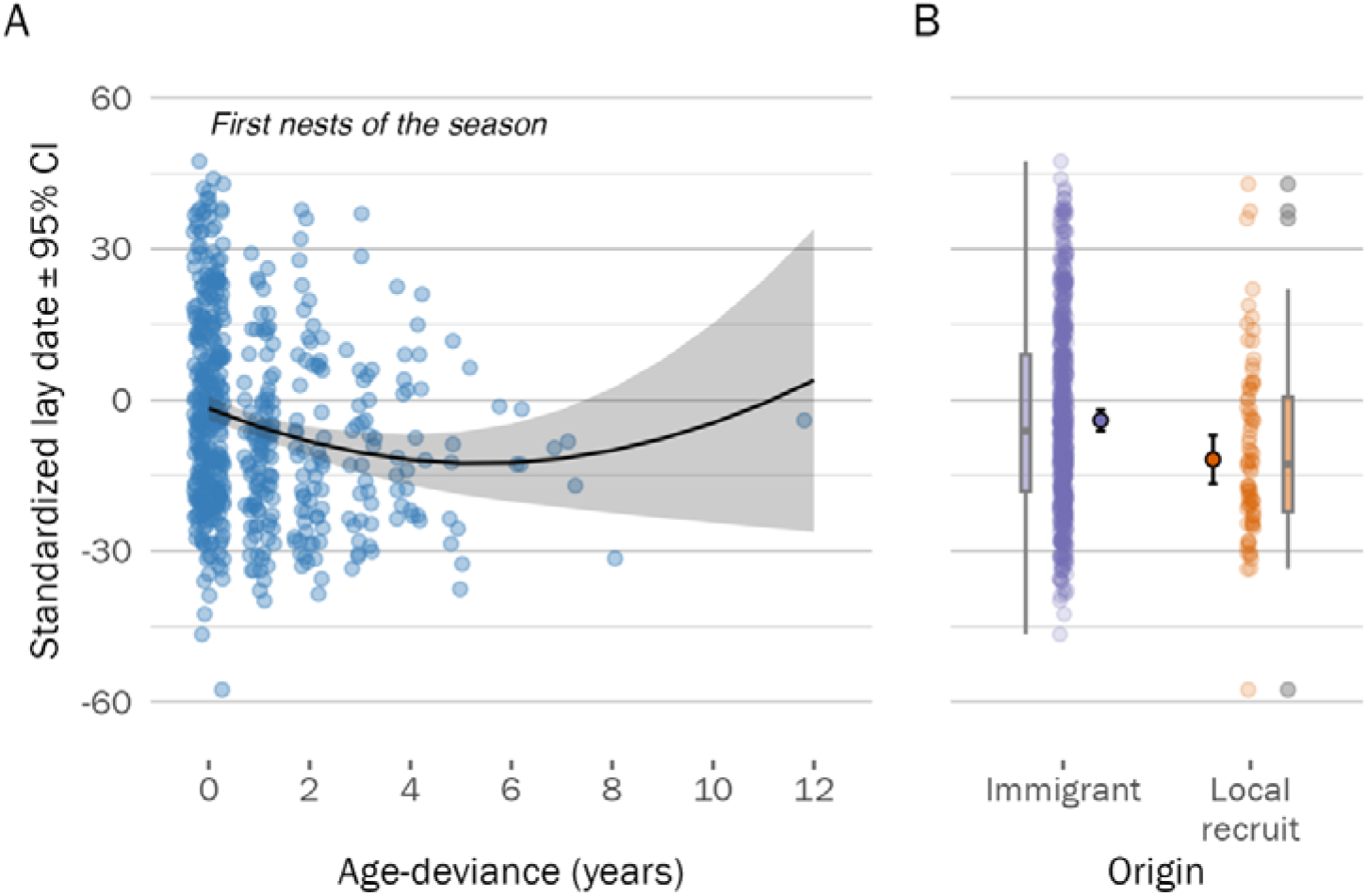
Age- and origin-dependent breeding phenology of female snowy plovers. A) Within-individual variation in age-specific nest initiation date – as females gained more experience in the local population, they started nesting earlier, however this trend reversed at older ages (albeit with high uncertainty). Each datum represents an individual’s ‘age-deviance’ (i.e., a within-group centered measure of the number of years since the individual’s first observed local breeding attempt, see *Methods* for more details) and the lay date of its first nest each year. B) Origin-specific variation in nest initiation date – females that hatched locally and recruited into the breeding population (orange) tended to nest earlier than birds originating from elsewhere (purple). Inner-most distributions show the model estimates and 95% CI, outer-most box plots show the inter-quartile ranges of the raw data (point-cloud).

**Table 3.** Sources of lay date variation. Note: within the *Partitioned R^2^* sub-table the term ‘Senescence’ describes the collective variation explained by the linear and quadratic within-individual age-deviance effects in the top sub-table, the term ‘Selective appearance’ and ‘Selective disappearance’ describe the variation explained by the between individual first- and last-breeding age fixed effects of the top sub-table, respectively.

| *Lay date of first nest* | *Mean estimate* | *95% confidence interval* |
| --- | --- | --- |
| **Fixed effects $\beta$ (standardized)** |  |  |
| Mother tarsus length | 0.02 | [-0.07, 0.1] |
| Local recruit | -7.80 | [-13.17, -3.05] |
| Within ind. age-deviance (linear) | -0.16 | [-0.26, -0.05] |
| Within ind. age-deviance (quadratic) | 0.09 | [0.01, 0.17] |
| Between ind. first breeding age | 0.06 | [-0.04, 0.16] |
| Between ind. last breeding age | -0.04 | [-0.18, 0.1] |
| **Partitioned $R^2$** |  |  |
| Total Marginal $R^2$ | 0.06 | [0.04, 0.12] |
| Total Conditional $R^2$ | 0.23 | [0.13, 0.38] |
| Mother tarsus length | 0.00 | [0, 0.06] |
| Local recruit | 0.02 | [0, 0.08] |
| Senescence | 0.02 | [0, 0.07] |
| Selective appearance | 0.01 | [0, 0.06] |
| Selective disappearance | 0.00 | [0, 0.06] |
| **Random effects $\sigma^2$** |  |  |
| Individual | 60.69 | [18.94, 117.67] |
| Year | 2.00 | [0, 12.89] |
| Residual | 283.38 | [229.95, 326.82] |
| **Adjusted repeatability $r$** |  |  |
| Individual | 0.19 | [0.06, 0.32] |
| Year | 0.01 | [0, 0.03] |
| Residual | 0.80 | [0.67, 0.93] |
| **Sample sizes $n$** |  |  |
| Years | 13 |  |
| Individuals | 376 |  |
| Observations (i.e., Nests) | 568 |  |

### Individual variation in egg volume

Overall, our model accounted for 74.4% of variation in egg volume, with fixed effects explaining 6.6% of this variation (Table 4). Females were highly repeatable in their egg volumes between clutches (*r* = 0.47; Table 4). Furthermore, eggs within the same clutch were moderately repeatable in volume (*r* = 0.16; Table 4). The strongest fixed effect explaining egg volume variation was the structural size of the mother (Fig. S3b, Table 4): larger females laid larger eggs than smaller females (model predicted difference in egg size between largest and smallest females in the sample: 0.57 *cm*^3^ [0.33, 0.80] 95% CI). The second strongest effect was the between-individual quadratic season function (Fig. 2c): eggs were smallest at the start of the season (model prediction: 7.18 *cm*^3^ [6.98, 7.38] 95% CI) and then increased, to be largest shortly after the middle of the season (model prediction: 7.68 *cm*^3^ [7.60, 7.75] 95% CI). Average egg volume also increased between sequential clutches within individuals but with smaller magnitude than the population-level trend (predicted increase of 0.20 *cm*^3^ [0.03, 0.37] 95%CI). Senescence in egg volume was not supported (Table 4). Furthermore, we found no support for selective (dis)appearance of individuals according to egg volume (Table 4).

**Table 4.** Sources of egg size variation. Note: within the *Partitioned R^2^* sub-table the term ‘Senescence’ describes the collective variation explained by the linear and quadratic within-individual age-deviance effects in the top sub-table, the term ‘Selective appearance’ and ‘Selective disappearance’ describe the variation explained by the between individual first- and last-breeding age fixed effects of the top sub-table, respectively; the term ‘Within ind. seasonality’ describes the variation explained by the within individual lay-date deviance effect in the top sub-table, and the term ‘Between ind. seasonality’ describes the collective variation explained by the linear and quadratic lay date effects in the top sub-table.

| **Egg volume** | **Mean estimate** | **95% confidence interval** |
| --- | --- | --- |
| **Fixed effects $\beta$ (standardized)** |  |  |
| Mother tarsus length | 0.22 | [0.14, 0.28] |
| Clutch size (< 3 eggs) | -0.03 | [-0.11, 0.04] |
| Within ind. age-deviance (linear) | -0.01 | [-0.08, 0.05] |
| Within ind. age-deviance (quadratic) | -0.04 | [-0.09, 0] |
| Between ind. first breeding age | 0.04 | [-0.04, 0.13] |
| Between ind. last breeding age | 0.03 | [-0.1, 0.15] |
| Within ind. lay-date-deviance | 0.09 | [0.05, 0.13] |
| Between ind. linear lay date | 0.16 | [0.1, 0.21] |
| Between ind. quadratic lay date | -0.06 | [-0.12, -0.01] |
| **Partitioned $R^2$** |  |  |
| Total Marginal $R^2$ | 0.07 | [0.05, 0.12] |
| Total Conditional $R^2$ | 0.66 | [0.63, 0.7] |
| Mother tarsus length | 0.04 | [0.02, 0.09] |
| Clutch size (< 3 eggs) | 0.00 | [0, 0.05] |
| Senescence | 0.00 | [0, 0.05] |
| Selective appearance | 0.00 | [0, 0.05] |
| Selective disappearance | 0.00 | [0, 0.05] |
| Within ind. seasonality | 0.01 | [0, 0.05] |
| Between ind. seasonality | 0.01 | [0, 0.06] |
| **Random effects $\sigma^2$** |  |  |
| Nest / Individual | 0.03 | [0.03, 0.04] |
| Individual | 0.10 | [0.08, 0.12] |
| Year | 0.01 | [0, 0.01] |
| Residual | 0.08 | [0.07, 0.08] |
| **Adjusted repeatability $r$** |  |  |
| Nest / Individual | 0.16 | [0.12, 0.2] |
| Individual | 0.45 | [0.39, 0.5] |
| Year | 0.03 | [0.01, 0.06] |
| Residual | 0.37 | [0.33, 0.41] |
| **Sample sizes $n$** |  |  |
| Years | 14 |  |
| Individuals | 424 |  |
| Nests | 836 |  |
| Observations (i.e., Eggs) | 2393 |  |

To further acknowledge uncertainty in the age-estimates provided by BaSTA, we re-analyzed the egg volume model using only known-aged females. Notably, this strengthened the effect sizes of ‘mother tarsus length’ and the within-individual age effects (i.e., ‘linear age’ and ‘quadratic age’). For example, the ‘mother tarsus length’ effect (β*_tarsus_*) increased to 0.32 [0.11, 0.50 95% CI], the ‘linear age’ effect (β*_age_*) increased to -0.05 [-0.21, 0.12], and the ‘quadratic age’ effect (β *^2^*) increased to -0.10 [-0.24, 0.05] (Table S2). However, given that the sample size of the known-aged female data set was greatly reduced, this analysis is arguably underpowered: the 95% confidence intervals bounding the effect sizes increased and, in many cases, overlapped zero. Furthermore, the bootstrap simulations incorporating the individual birth-year posteriors estimated from BaSTA confirmed the results of all age-dependent models (Fig. S4).

## DISCUSSION

Increased competition for mating opportunities in species with conventional sex roles is typically seen as a consequence of anisogamy where the production of many small gametes allows polygamous males to maximize their reproductive potential (Darwin, 1871; Bateman, 1948; Wade & Shuster, 2002; Schärer *et al*., 2012; Lehtonen *et al*., 2016). Polyandry, whereby females compete more intensively for mates than males, could lead to the evolution of smaller eggs in a similar way. Andersson (2004) developed a theoretical quantitative genetic model of female sexual selection by scramble competition over males in an attempt to explain reduced egg size in polyandrous species. Andersson’s (2004) model predicted that scramble competition for males should lead to egg size reduction and/or early breeding in sequentially polyandrous females of species with largely invariable clutch sizes. We tested the predictions of Andersson’s (2004) model empirically at the individual level: examining the links between breeding phenology, individual mating strategies, and egg size variation in a population of snowy plovers that contains monogamous and polyandrous females (Cruz-López *et al*., 2017; Eberhart-Phillips *et al*., 2017; Kupán *et al*., 2021). Consistent with Andersson’s (2004) predictions, we found that female polyandry was associated with early breeding and that early clutches typically contained smaller eggs than clutches laid later in the season – both between and within individual females. However, we interpret this pattern differently than Andersson (2004). In our population, early breeding females whose nesting attempts were successful invariably deserted the brood shortly after hatching (Cruz-López *et al*., 2017; Kupán *et al*., 2021) resulting in collinearity between mating behavior and first lay date. We considered lay date therefore as a proxy for polyandry potential, with early nesting females having a higher probability to desert their offspring and become polyandrous (unless their first nest got predated or failed).

Although the statistical associations between breeding phenology, egg size, and female mating behavior are intriguing, we note that their impact on the egg size variation observed in this population is rather limited. First, despite snowy plovers’ unusual mating system with sequential polyandry, monogamy, and occasionally polygyny occurring in the same population (Warriner *et al*., 1986; Eberhart-Phillips *et al*., 2017), the egg size variation we report is not more pronounced than in other birds: the proportional difference between the largest and smallest egg in our population was approximately 50%, which is of very similar magnitude to the intraspecific range observed within other bird populations (Christians, 2002). Other constraints may therefore provide limitations on egg size variation in snowy plovers. Waders produce relatively large eggs that provide vital reserves for their precocial young and the moderate egg size variation observed in snowy plovers could indicate that females are unable to reduce egg size further without jeopardizing chick viability. Second, we note that structural size of females had a stronger effect on egg size variation than breeding phenology. This is consistent with other studies (Christians, 2002; Giraldo-Deck *et al*., 2022) showing that larger females produce larger eggs than smaller females. Third, we found that, as demonstrated in other studies (e.g., Christians, 2002), female identity explained the largest part of the variation in egg size, with repeatability of egg size in females was very high. High repeatability suggests that egg size may have considerable genetic variation, although it has been suggested that many loci with small effects seem to encode this life history trait (Santure *et al*., 2013).

In contrast to egg size, female mating behavior showed a very low individual repeatability across years. The observed low individual consistency in polyandry is most likely related to the highly stochastic nature of nest survival, where the main sources of breeding failure in our population are flooding, predation, and chick mortality through starvation (Cruz-López *et al*., 2017). Early breeding females whose nests failed, typically stayed together with their mate for rapid re-nesting (Halimubieke *et al*., 2019). These females may have intended to become polygamous but since their replacement clutch hatched later in the season when conditions for chicks or remating are less favorable, they switched to a monogamous strategy that involves longer brood care (Kupán *et al*., 2021). This was supported by the higher repeatability multi-clutching behavior (i.e., pooling polyandrous and re-nesting females). Snowy plover females can adjust their brood care efforts flexibly according to the value and needs of the young (Kupán *et al*., 2021). Female care and re-mating efforts are inversely related to each other and show strong seasonality. Early in the breeding season when survival prospects for the chicks are favorable, re-mating opportunities are high, and their mate can successfully rear the chicks alone, females typically desert their broods right after hatching of the chicks to quickly start a new breeding attempt with another male (Székely *et al*., 1999; Cruz-López *et al*., 2017; Kupán *et al*., 2021). Females whose chicks hatch later in the season, typically remain with the brood and provide care jointly with their mate. However, desertion during brood care is tightly associated with chick mortality (Kupán *et al*., 2021). When survival prospects of their offspring are deteriorating and chicks die despite the joint care, initially caring females desert their broods and presumably remate elsewhere where environmental conditions are better. Such high plasticity in female care and mating strategies allows females to flexibly respond to nest and chick survival, and is an important ecological modulator of breeding behavior in snowy plovers. As expected, in years that females could be polyandrous, they produced more chicks than in years when they were monogamous, which we attribute to 1) these females laying early enough to allow time for multiple breeding attempts, and 2) having good fortune that their first nest was not depredated, so that they could desert and pursue a sequential mate.

Most studies conducted on temperate or high latitude breeding shorebirds have found a negative association between time of the season and egg size (Byrkjedal & Kalas, 1985; Sandercock *et al*., 1999; Kubelka *et al*., 2020) although in polyandrous red-necked phalaropes (*Phalaropus lobatus*) egg size increased throughout the breeding season (Kwon et al., 2018, n.b., the presented effect size is small). However, many investigations of seasonal egg size dynamics did not disentangle whether the observed changes were due to within- or between-individual effects. A study that included 15 arctic shorebirds suggested that between-individual variation may account for more of the seasonal variation in egg size than within-individual variation (Weiser *et al*., 2018). In our population, early season nesters produced smaller eggs on average than females initiating nests at a later date. These seasonal changes were driven by within individual changes: females generally increased egg volume between consecutive nesting attempts (albeit the effect size was small), suggesting that maternal investment during early breeding attempts could be impaired by physiological limitations. Mutually non-exclusive constraints may include low food availability (Steigerwald *et al*., 2015), and carry-over costs associated with intense competition (Duckworth, 2006) or migratory status (Crossin *et al*., 2010). In contrast, delayed breeders can take advantage of a slower pace to gather more resources and maximize investment prior to clutch initiation – whilst having only enough time for a single breeding attempt. Although we did not collect any direct empirical data on temporal changes in food availability during our study (e.g., with pit-fall traps), the invertebrate prey communities of salina ecosystems (i.e., like that of our study population) are well documented to have dynamic emergence and quiescent phenologies (Britton & Johnson, 1987), providing important ecological context and support of our notion that resource availability for egg-laying female snowy plovers is highly seasonal. The observed decrease of egg size in clutches produced at the end of season is consistent with such constraints for breeding females as females are in a rush to complete their clutches and finish incubation before the onset of catastrophic floods caused by increased precipitation and tide height at the end of the breeding season (Plaschke *et al*., 2019).

Despite being long-lived – the oldest documented female in our sample was at least 13 years old (species’ longevity record: 20 years; Colwell et al., 2017; M. A. Colwell *personal communication,* 15 March, 2021) – and investing substantially in reproduction year-after-year, we found no evidence of age-dependent trade-offs in egg size or polyandry potential. Contrary to our predictions derived from senescence theory, older females initiated nests earlier in the season compared to their younger conspecifics – indicating age-dependent competitive ability that may initially increase the polygamy potential of females (Oring *et al*., 1994). Age-dependent variation in lay date followed a non-linear pattern: lay date advanced with each year of age until a peak at age six, after which the age-lay date trajectory was unclear because of limited sampling in older age classes. In snowy plovers, occasionally females re-mate with or retain their last mate in the following season (Halimubieke *et al*., 2019), presumably to skip the time consuming mate search and start breeding early – a process that could also explain the age-dependent advance in lay date we observed. Furthermore, locally recruited females (i.e., hatched locally) bred earlier than immigrant females indicating a role for prior experience giving competitive advantage for breeders in our population.

Egg size was not clearly associated with female age. Several studies of oviparous organisms have observed age-dependent variation in egg size, with some studies finding a positive relationship (Cooch *et al*., 1992; Flint & Sedinger, 1992; Robertson *et al*., 1994; Warner *et al*., 2016; Verhoeven *et al*., 2020) and others observing a negative relationship (Reid, 1988; Potti, 1993; Ito, 1997). Recent longitudinal studies of wild avian systems showed a quadratic relationship with a prominent increase in egg size in early life, followed by a peak and then a late-life decline (Bouwhuis *et al*., 2009; Jankowiak *et al*., 2018). In captive avian systems, the early life increase in egg size is also observed, but not the late-life decline likely due to *ad libitum* food (Woodard & Abplanalp, 1971). In our study, a quadratic-age model of senescence in egg size was not statistically clear, and did not change when we repeated the analysis including only the 45 females of known age (10% of the original sample). We acknowledge that our relatively small sample of known-age individuals presents a limitation to our study – we cannot exclude a moderate effect of senescence on egg size. Nevertheless, potential explanations for the lack of a clear senescence trend observed in egg size are that 1) our open- and free-living population lacks extremely old individuals that would exhibit the intrinsic “costs paid” to somatic loss, and/or 2) there is little room for further reduction because of viability constraints for the chicks. In shorebirds, chicks are not fed by the parents but rather must forage for themselves immediately after hatching and larger chicks that hatch from larger eggs typically survive better than smaller chicks (Blomqvist *et al*., 1997; Giraldo-Deck *et al*., 2022). The extra nutrients provided by a large egg may be crucial for the chick survival during the first few days of life when foraging efficiency is reduced as chicks still learn how to catch prey (Ricklefs, 1968). Consequently, additional reductions of egg size may not be feasible as the knock-on effects for chick survival are too costly (Williams, 1994; Starck & Ricklefs, 1998), therefore limiting a females’ potential to save resources by reducing egg volume. Chick-viability constraints may also provide an explanation why the variation in egg size at the population level is relatively low.

### Conclusions

Past species-level studies have linked polyandry and sex-role reversal to reduced female gamete size (Slotow, 1996; Andersson, 2004), as smaller eggs would permit females to produce several clutches in rapid succession (Liker *et al*., 2001). In our individual-level investigation we found that early nesting females were more likely to be polyandrous and produced clutches containing smaller eggs than females that started their first breeding attempt later in the season. Starting to breed early in the season is an important pre-determinant of female polygamy. Prior experience can provide both older and local females a competitive advantage over younger and naïve conspecifics scrambling for early breeding opportunities that are at the heart of the polyandrous mating strategy (Andersson, 2004). However, we found no evidence supporting an adaptive egg-size adjustment to obtain a competitive advantage in the scramble competition for mates. Rather we suspect that the observed seasonal variation in egg size can also be explained by other constraints that include a survival advantage for early hatching chicks, resource constraints during the egg-producing phase and, most importantly, between individual differences. Senescence did not clearly have a major impact on egg size variation nor on the mating behavior of females. Future research of polyandrous systems should also examine the impact of seasonality, age, and mating behavior on the reproductive fitness of males, as males provide a larger share of parental care than females in these systems and therefore should have a more direct impact on the survival prospects of the young. Examining the effects of season- and age-dependent trade-offs on individual fitness and variation in mating tactics will enable a deeper understanding about selective pressures and constraints of diverse and plastic sex roles in natural populations.

## DATA ACCESSIBILITY

We provide all computer code and documentation as a html vignette written in Rmarkdown (click here: <u>File S1</u>) together with all the raw datasets needed to reproduce our modeling and analyses – these files can be found in this project’s Open Science Framework repository: <u>doi.org/10.17605/OSF.IO/UCW6J</u>

## SUPPLEMENTARY TABLES

**Table S1.** Relationship between the average egg volume of a clutch and the average chick weight at hatching. See Supplementary Text D for methods.

| *Chick hatch weight* | *Mean estimate* | *95% confidence interval* |
| --- | --- | --- |
| Fixed effects $\beta$ (logit standardized) | | |
| Average egg volume of clutch | 0.60 | [0.55, 0.66] |
| Partitioned $R^2$ | | |
| Total Marginal $R^2$ | 0.37 | [0.3, 0.43] |
| Total Conditional $R^2$ | 0.47 | [0.38, 0.56] |
| Random effects $\sigma^2$ | | |
| Mother identity | 0.01 | [0, 0.02] |
| Year | 0.01 | [0, 0.02] |
| Residual | 0.09 | [0.08, 0.11] |
| Adjusted repeatability $r$ | | |
| Mother identity | 0.05 | [0, 0.12] |
| Year | 0.05 | [0.01, 0.11] |
| Residual | 0.90 | [0.81, 0.98] |
| Sample sizes $n$ | | |
| Year | 14 |  |
| Individuals | 426 |  |
| Observations (i.e., Nests) | 841 |  |

**Table S2.** Sources of egg volume variation when using only known-aged females. Note: within the *Partitioned R^2^* sub-table the term ‘Senescence’ describes the collective variation explained by the linear and quadratic within-individual age effects in the top sub-table, the term ‘Selective appearance’ and ‘Selective disappearance’ describe the variation explained by the between individual first- and last-breeding age fixed effects of the top sub-table, respectively; the term ‘Within ind. seasonality’ describes the variation explained by the within individual lay date effect in the top sub-table, and the term ‘Between ind. seasonality’ describes the collective variation explained by the linear and quadratic lay date effects in the top sub-table.

| ***Egg volume (known-age subset)*** | ***Mean estimate*** | ***95% confidence interval*** |
| --- | --- | --- |
| **Fixed effects $\beta$ (logit standardized)** |  |  |
| Mother tarsus length | 0.32 | [0.11, 0.5] |
| Clutch size (< 3 eggs) | -0.02 | [-0.22, 0.19] |
| Within ind. age-deviance (linear) | -0.05 | [-0.21, 0.12] |
| Within ind. age-deviance (quadratic) | -0.10 | [-0.24, 0.05] |
| Between ind. first breeding age | 0.03 | [-0.31, 0.35] |
| Between ind. last breeding age | -0.01 | [-0.38, 0.37] |
| Within ind. lay-date-deviance | 0.06 | [-0.09, 0.2] |
| Between ind. linear lay date | 0.14 | [-0.02, 0.31] |
| Between ind. quadratic lay date | -0.16 | [-0.32, 0.01] |
| **Partitioned $R^2$** |  |  |
| Total Marginal $R^2$ | 0.12 | [0.06, 0.32] |
| Total Conditional $R^2$ | 0.66 | [0.58, 0.77] |
| Mother tarsus length | 0.08 | [0.02, 0.28] |
| Clutch size (< 3 eggs) | 0.00 | [0, 0.22] |
| Senescence | 0.01 | [0, 0.23] |
| Selective appearance | 0.00 | [0, 0.22] |
| Selective disappearance | 0.00 | [0, 0.21] |
| Within ind. seasonality | 0.04 | [0, 0.25] |
| Between ind. seasonality | 0.00 | [0, 0.22] |
| **Random effects $\sigma^2$** |  |  |
| Nest / Individual | 0.05 | [0.03, 0.08] |
| Individual | 0.06 | [0.02, 0.12] |
| Year | 0.00 | [0, 0.02] |
| Residual | 0.07 | [0.06, 0.09] |
| **Adjusted repeatability $r$** |  |  |
| Nest / Individual | 0.28 | [0.13, 0.43] |
| Individual | 0.33 | [0.12, 0.51] |
| Year | 0.01 | [0, 0.08] |
| Residual | 0.38 | [0.28, 0.51] |
| **Sample sizes $n$** |  |  |
| Years | 14 |  |
| Individuals | 43 |  |
| Nests | 112 |  |
| Observations (i.e., Eggs) | 320 |  |

**Table S3.** Relationship between the lay date of an individual’s first nest and their likelihood of re-nesting or being polyandrous (i.e., “multi-clutching”) in a given year. Fixed effect size of ‘first nest lay date’ is the standardized estimate on the logit scale.

| *Multi-clutching probability* | *Mean estimate* | *95% confidence interval* |
| --- | --- | --- |
| Fixed effects $\beta$ (logit standardized) | | |
| First nest lay date | -1.83 | [-2.3, -1.52] |
| Age since first breeding | -0.04 | [-0.26, 0.19] |
| First breeding age | 0.11 | [-0.12, 0.33] |
| Female tarsus length | -0.03 | [-0.25, 0.19] |
| Partitioned $R^2$ | | |
| Total Marginal $R^2$ | 0.37 | [0.28, 0.47] |
| Total Conditional $R^2$ | 0.40 | [0.3, 0.51] |
| First nest lay date | 0.37 | [0.28, 0.46] |
| Age since first breeding | 0.00 | [0, 0.17] |
| First breeding age | 0.00 | [0, 0.17] |
| Female tarsus length | 0.00 | [0, 0.17] |
| Random effects $\sigma^2$ | | |
| Individual | 0.35 | [0, 0.86] |
| Year | 0.40 | [0, 0.69] |
| Adjusted repeatability $r$ | | |
| Individual | 0.02 | [0, 0.12] |
| Year | 0.03 | [0, 0.08] |
| Sample sizes $n$ | | |
| Years | 14 |  |
| Individuals | 426 |  |
| Observations (i.e., Nests) | 665 |  |

## SUPPLEMENTARY FIGURES

**Figure S1.**
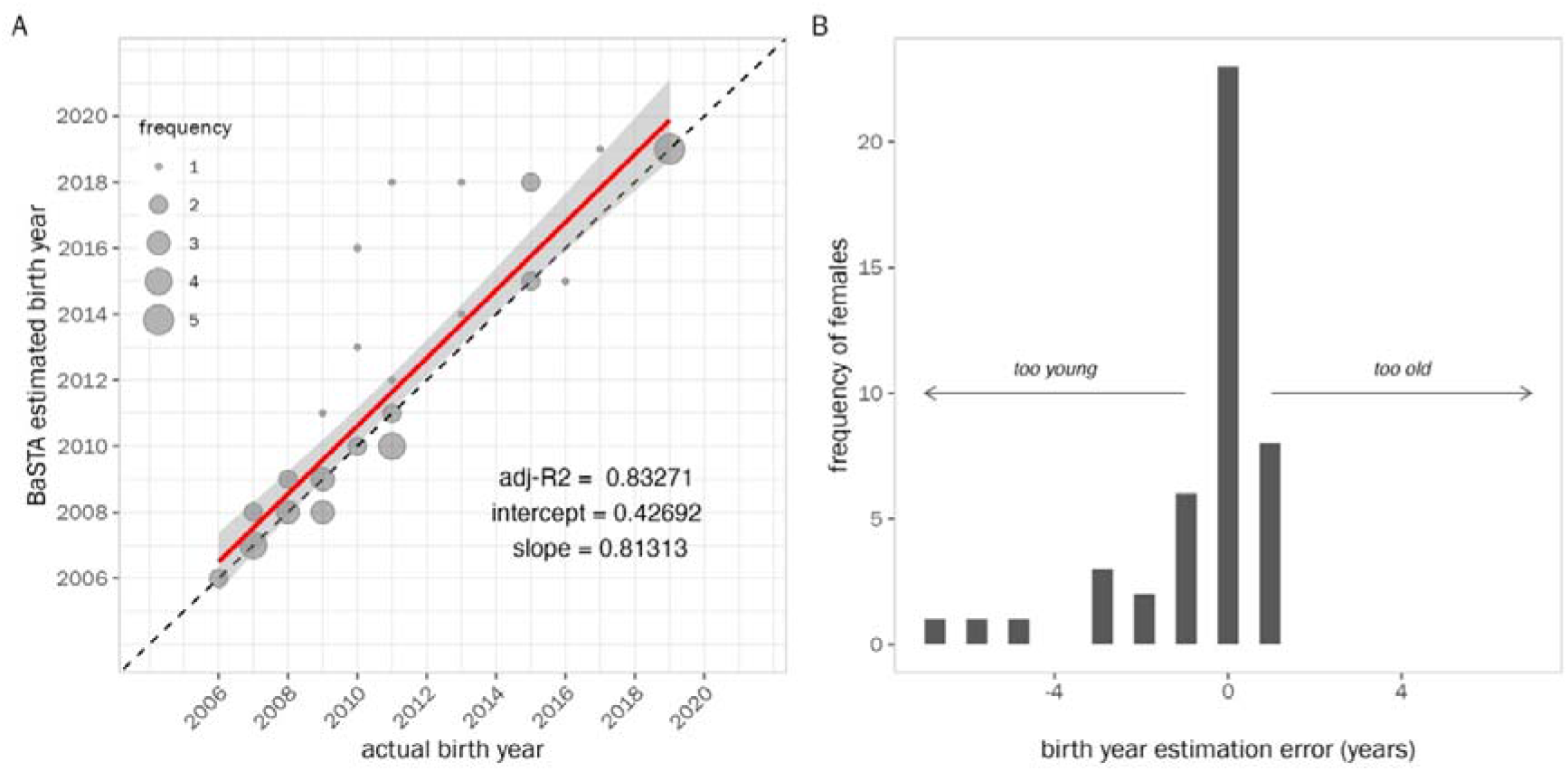
Results of simulation to cross-validate the birth year estimates of known-aged females produced by BaSTA. Panel A) shows a correlation between the estimated BaSTA birth year and the true birth year (size of points refers to the frequency of females), with the dashed diagonal illustrating a slope of 1 (i.e., a perfect estimation). Panel B) shows the distribution of the birth year error for all known-aged females based on the cross-validation simulation.

**Figure S2.**
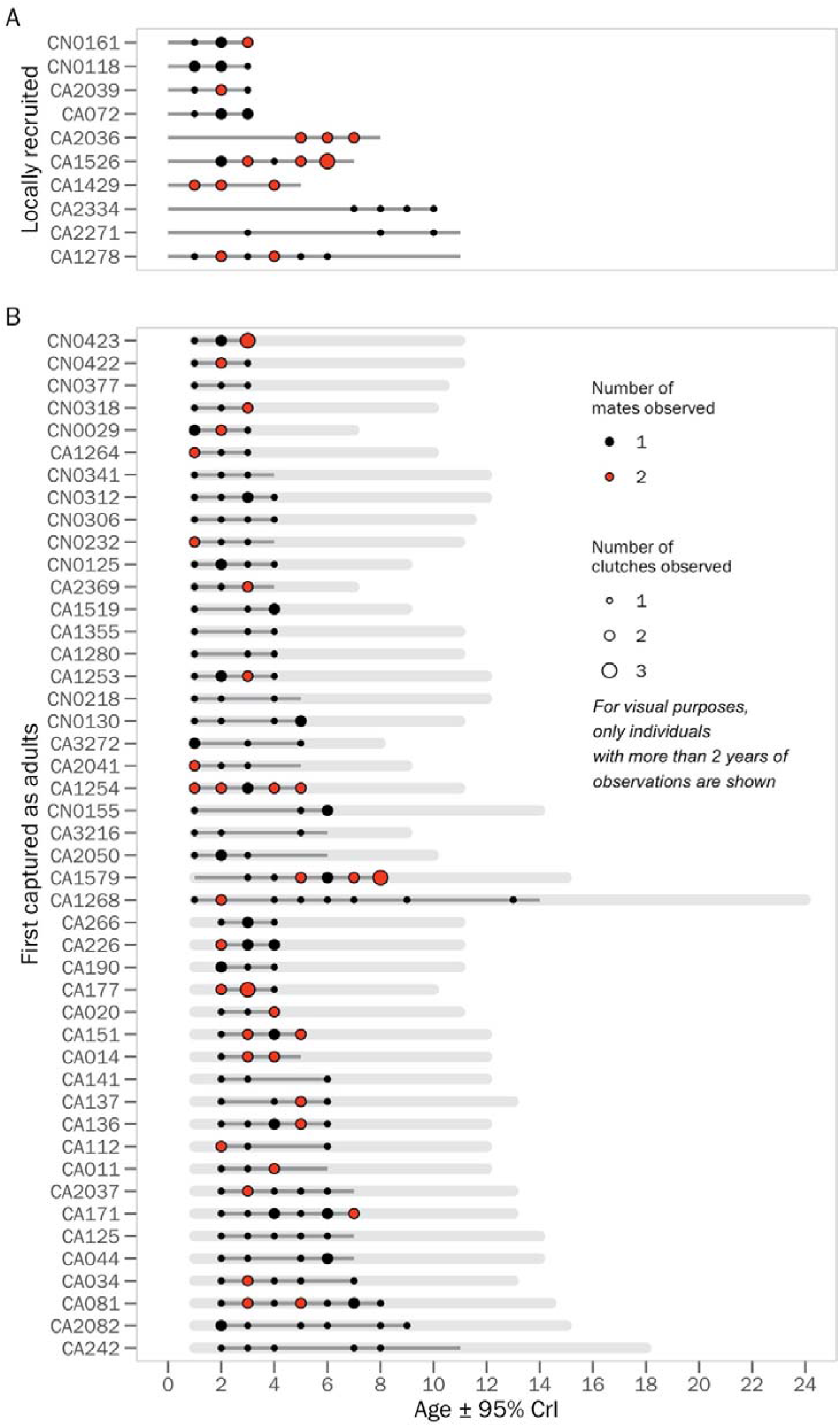
Mating strategy and clutch number of female snowy plovers according to age. Each row shows an individual female in the population for which we have at least three years of observations (note that our analysis also includes females with one or two years of observation, but given space constraints only individuals with a minimum of three years are plotted in this graph). Panel A) shows known-aged females which were born locally, whereas B) shows females that were initially captured as adults and are therefore of unknown age. Points illustrate the age at which we collected observations of egg volume, with the size of the point corresponding to the number of clutches measured at a given age, and the colour indicating if we observed the female mating with one or two distinct males (i.e., in case of multiple clutches at a given age). The light grey buffer around unknown-age females indicates the 95% CrI of the ages for an individuals’ observed period (i.e., lower limit indicates the minimum age the individual could have entered the population and the upper limit indicates the maximum age of an individual’s last observation based on BaSTA’s birth year posterior). The dark grey lines indicate the period for which an individual was observed alive (i.e., in some cases we encountered an individual in the field and confirmed its survival, but we did not observe its nest. Note also that the age at first encounter of all known-aged individuals is 0).

**Figure S2.**
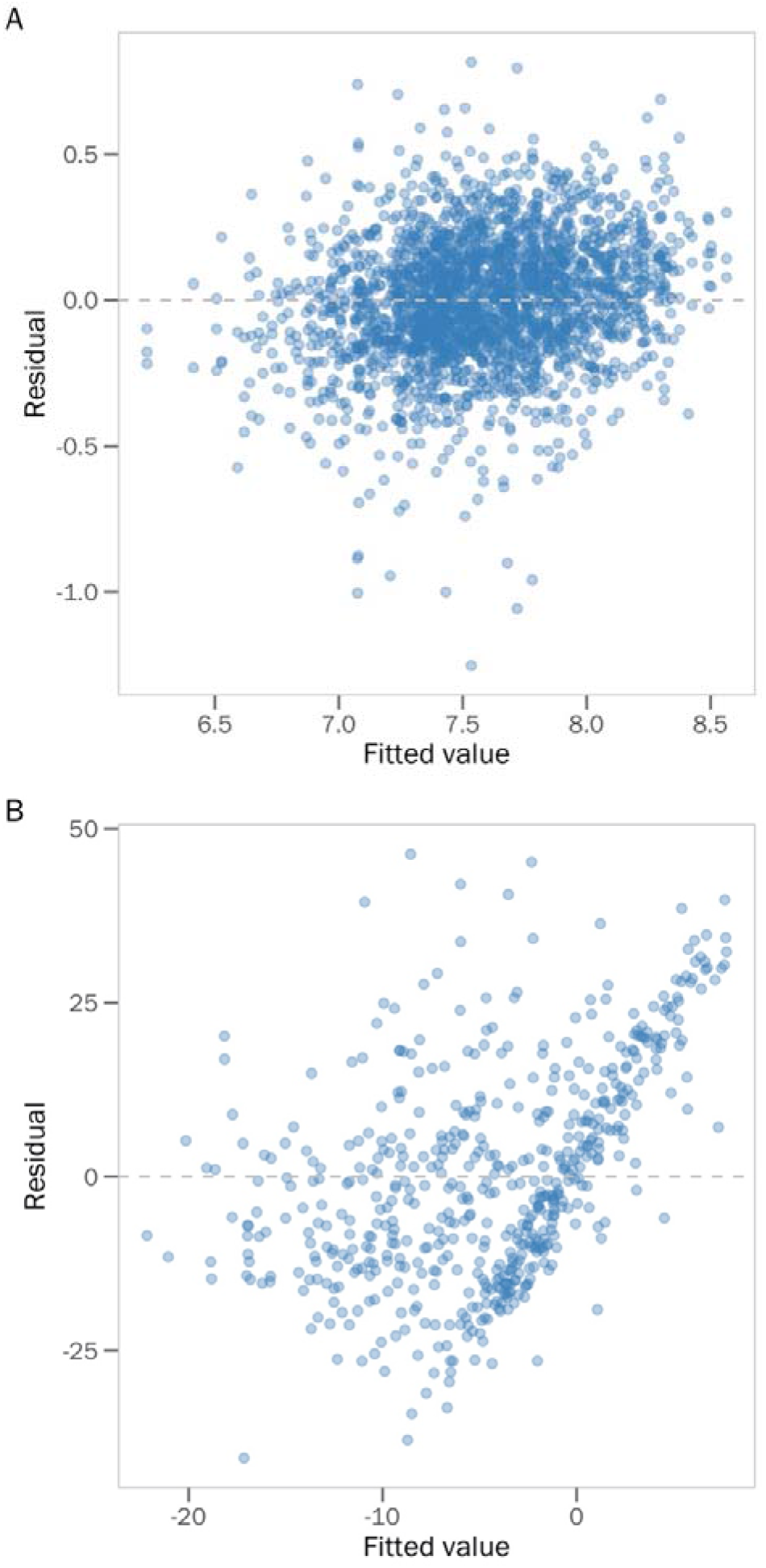
Homoscedastic spread of residuals from the A) ‘egg volume’ and B) ‘lay date’ models.

**Figure S3.**
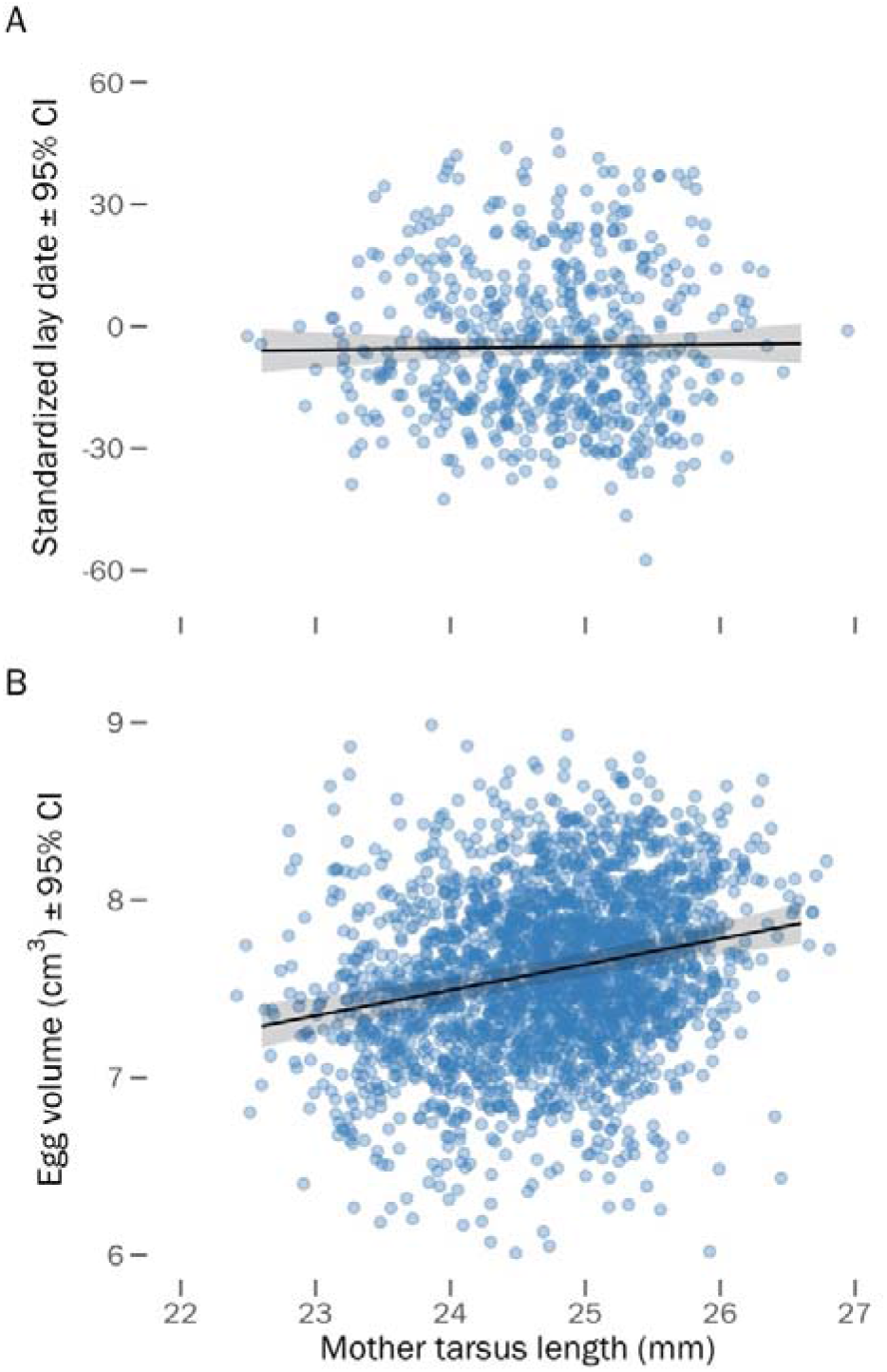
Relationship between a female’s structural size (i.e., her tarsus length) and A) the lay date of her first nest of the season, and B) the volume of her eggs.

**Figure S4.**
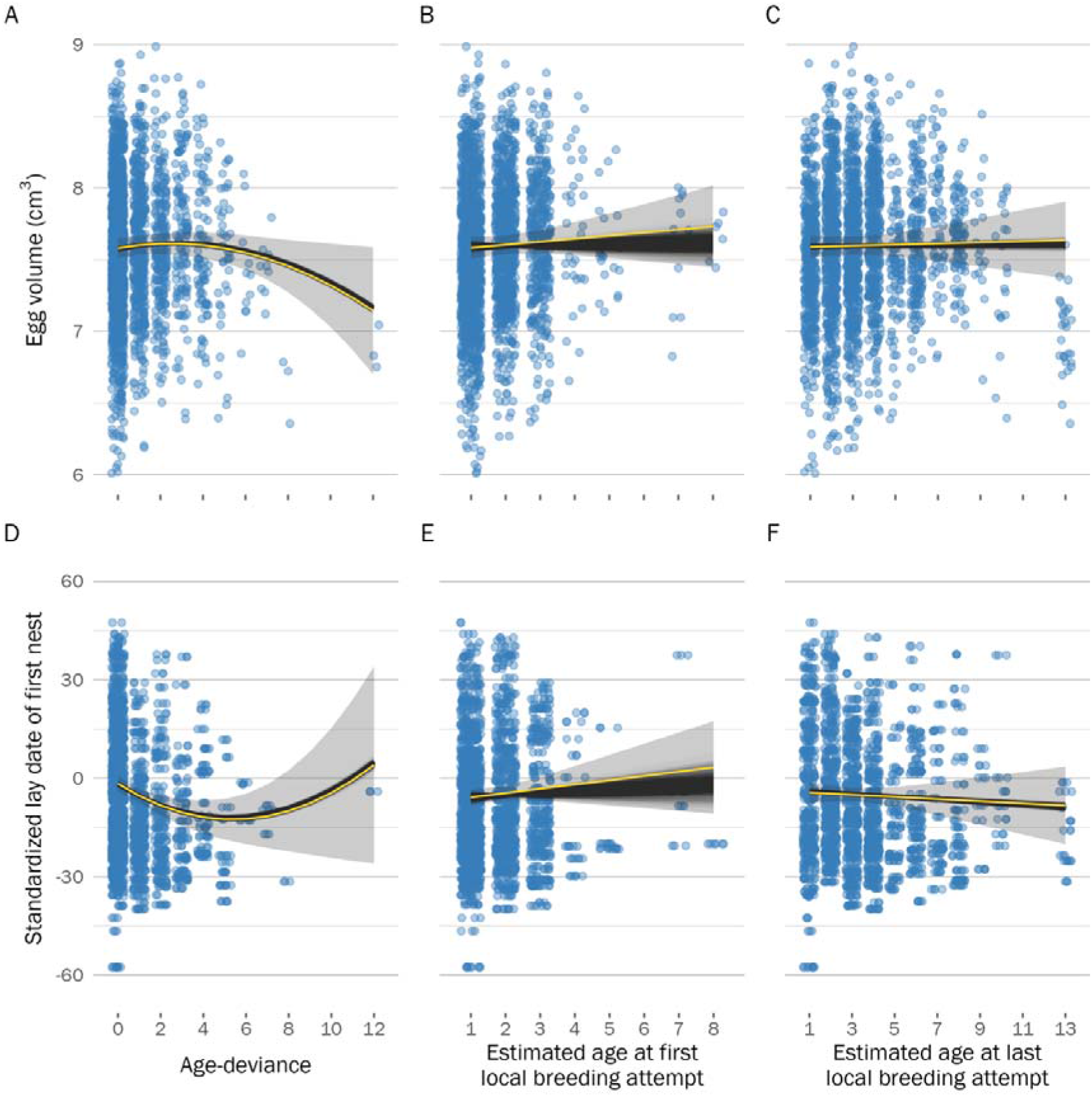
Visualization of the effect of uncertainty in the age estimate provided by BaSTA on age-dependent trends in egg volume (top row) and lay date (bottom row) dynamics. Black trends show the model predictions for the 1000 bootstraps of the BaSTA age estimate posteriors (see *Methods*). Panels A and D show the within-individual trends of the ‘age-deviation’ score – as expected, these measures are not impacted by uncertainty in the BaSTA age estimate because they are centered for each individual (i.e., the absolute age is irrelevant). Panels B and E show the between-individual trend of the ‘age at first breeding’ (i.e., selective appearance), and panels C and F show the between-individual trend of the ‘age at last breeding’ (i.e., selective disappearance). Yellow trends and grey ribbons visualize the 95% CI of the model predictions when using the median age estimate provided by BaSTA (i.e., the effect sizes of the ‘egg volume’ and ‘lay date’ models shown in Fig. 1).

**Figure S5.**
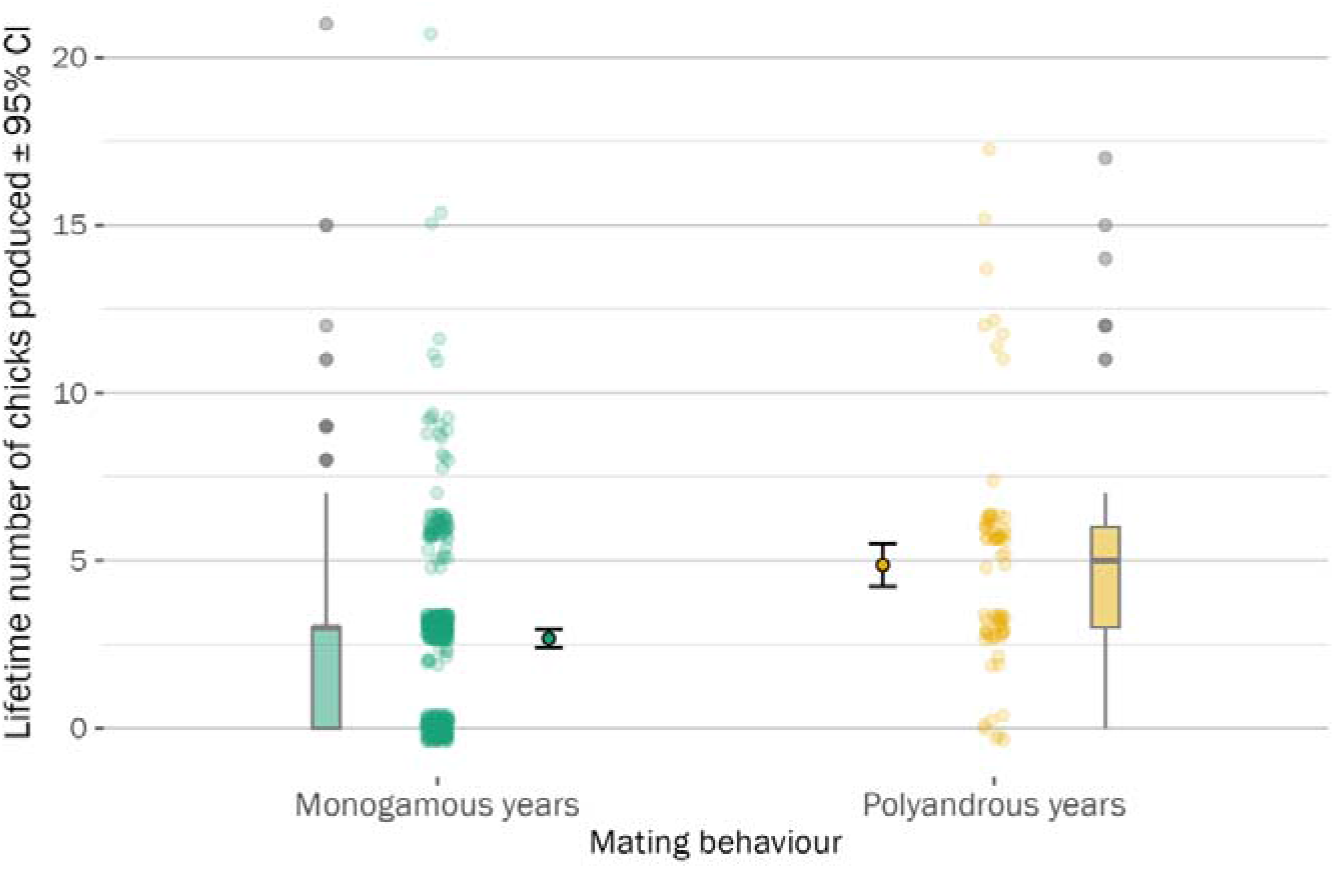
The total observed number of chicks produced by females in years when they were polyandrous was 2.18 (1.83–2.55 95% CI) chicks more than in years when were monogamous (Fig. S9; *N* = 14 years, 424 females, 836 nests).

## Notes

https://leberhartphillips.github.io/snowy_plover_eggs/

https://doi.org/10.17605/OSF.IO/UCW6J

### Competing Interest Statement

The authors have declared no competing interest.

### Summary of Updates

Updated version of the manuscript after addressing reviewers comments.

