## Supplementary Materials for "Egg size variation in the context of polyandry: a case study using long-term field data from snowy plovers"

**SUPPLEMENTARY TEXTS**

**Supplementary Text A**

*Age estimation of individuals with unknown origin*—Investigating age-dependent processes in the wild is challenging, as analyses often involve a mix of individuals of known or unknown age (Colchero *et al.*, 2012) – with the former being initially marked at birth (i.e., uncensored), and the latter being immigrants of unknown age or those that were born before the study’s first marking occasion (i.e., left-truncated). To estimate the ages of unknown individuals in our marked population, we employed a capture-mark-recapture analysis using the ‘Bayesian Survival Trajectory Analysis’ (BaSTA) package in R (v1.9.4, Colchero *et al.*, 2012), which uses a Bayesian hierarchical framework to fit parametric survival functions of the marked population while accounting for imperfect detection. Furthermore, BaSTA derives estimates of birth year of left-truncated individuals from the population mean of the projected survival function. As snowy plovers show sex differences in survival (Eberhart-Phillips *et al.*, 2017; 2018), we used female-specific survival functions for this study. Due to high natal dispersal, we could not confidently determine the fate of juveniles marked in our population. To acknowledge this uncertainty, our capture-mark-recapture sample only included individuals that survived their first breeding season, i.e., we constrained first-year survival probability to 1.

In total, our capture-mark-recapture data comprised records of 450 uniquely marked females, of which 45 hatched locally and subsequently recruited into the adult population as known-age individuals (Fig. S2a), and the remaining 405 females were adults of unknown age and origin. We monitored the presence or absence of marked individuals by recapturing or observing them in the field in all study years except for 2014, amounting to a total of 916 post-birth annual detections of the 450 females in our sample (median annual detections per adult = 2; mean = 2.04, 1.45 SD). We discarded annual detections if a bird was only encountered once via a resighting of a color combination, as single-observations are known to influence capture-mark-recapture analyses (Tucker et al. 2019). A logistic bathtub-shaped mortality model had the best fit to our data – revealing that female mortality rate increased until age 5 years, after which it became constant (Fig. S2b; see Appendix S1 for detailed methods). Using this model, we extracted the birth year estimate posteriors for each unknown-age individual in the capture-mark-recapture sample. Note that three individuals (one first encountered as an adult [CA1579] and two locally hatched recruits [CA2036 and CA1526]; Fig. S3) had been already marked two years prior to the start of our monitoring period (i.e., pre-2006) and were thus added to our sample after running BaSTA on capture-mark-recapture data for 2006–2020.

To explain how mortality risk is associated with age, we compared four commonly used survival functions: exponential, Gompertz, logistic, and Weibull models. The exponential function keeps survival constant across age

(Cox & Oakes, 1984), whereas the other functions allow for age-dependent variation in survival: the Gompertz model is an exponential function in which age-specific mortality is scaled by baseline mortality (Gompertz, 1812; Pletcher, 1999) and the Weibull model is a power function which assumes that baseline and age-specific mortality rates are independent (Pinder *et al.*, 1978). In addition to the stand-alone versions of these functions, we considered two alternative forms of the Gompertz, logistic, and Weibull functions – “Makeham” and “bathtub” shapes. The Makeham shape constrains the survival function to converge to a constant, rather than zero, as age increases (Pletcher, 1999), whereas the bathtub shape enables concavity in the function, such that mortality could decrease at early ages, but increase later in life (Siler, 1979).

We used four parallel simulations to run the Markov chain Monte Carlo (MCMC) optimization procedure in BaSTA with 800,000 iterations, a 100,000 burn-in period, and a thinning interval of 2000 to minimize serial autocorrelation in the chain (see Fig. S2 for simulation diagnostics). We ranked the survival models according to their deviance information criterion (*DIC*) and determined that the logistic model with a bathtub shape fitted our data best (i.e., lowest *DIC*, (Spiegelhalter *et al.*, 2002); Table S1). This logistic model revealed that female mortality rate increased until age 5 years, after which it became constant (Fig. 1b). Using the top model, we extracted the point estimate and 95% credible interval birth year for each individual in the capture-mark-recapture sample.

Tucker, A.M., McGowan, C.P., Robinson, R.A., Clark, J.A., Lyons, J.E., DeRose-Wilson, A., Du Feu, R., Austin, G.E., Atkinson, P.W. & Clark, N.A., Effects of individual misidentification on estimates of survival in long-term mark–resight studies, *The Condor*, **1211**, duy017

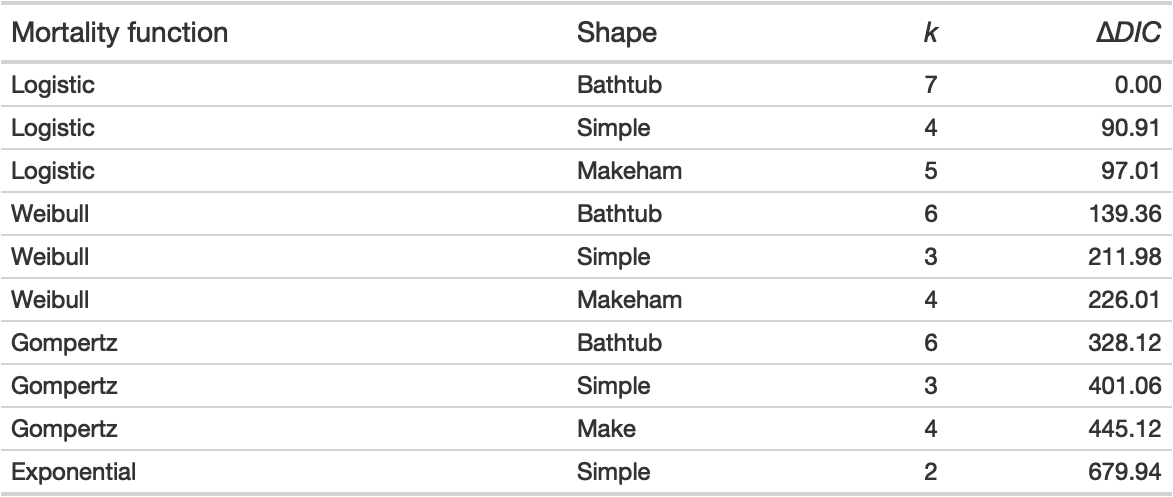
**Table S-A1.** Hierarchical capture-mark-recapture models used to describe mortality patterns of snowy plovers at Bahía de Ceuta, Mexico, between 2006 and 2020. *k:* number of modelled parameters; Δ*DIC:* difference in deviance information criterion (*DIC*) between a given model and the top model. *DIC* value of top model was 5279.1.

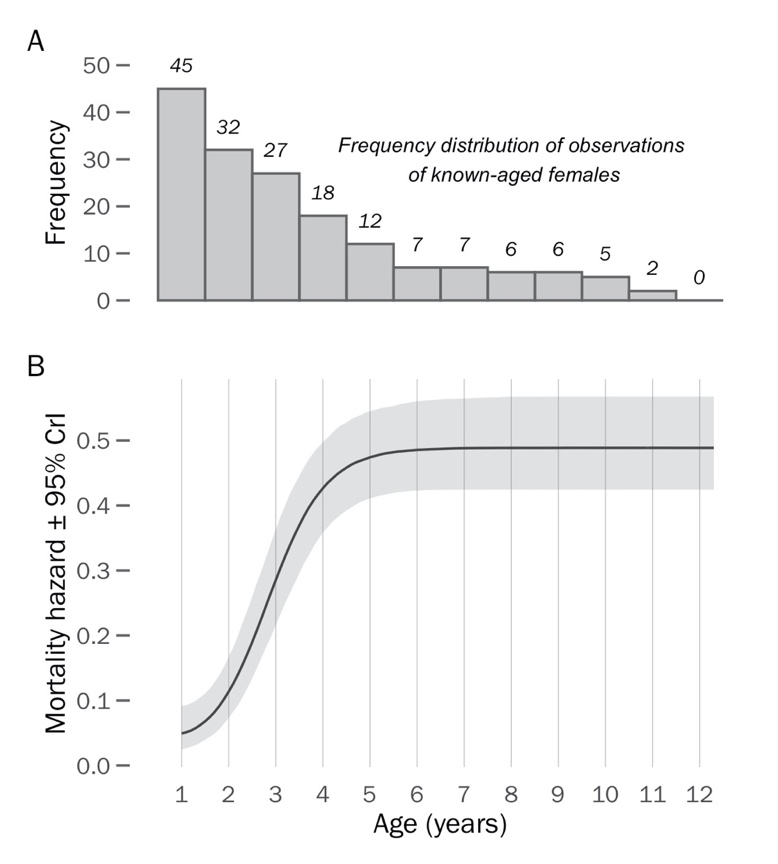

**Figure S-A1.** Logistic-bathtub mortality function for snowy plover females: a) frequency distribution of age-specific observations of 45 known-aged females and b) age-dependent mortality hazard.

**Figure S-A2.** Bayesian model diagnostics of the Logistic bathtub shaped survival model. a) Serial autocorrelation of iterations within chains, b) chain convergence (*R-hat*) of model, and c) trace plots of chain dynamics.

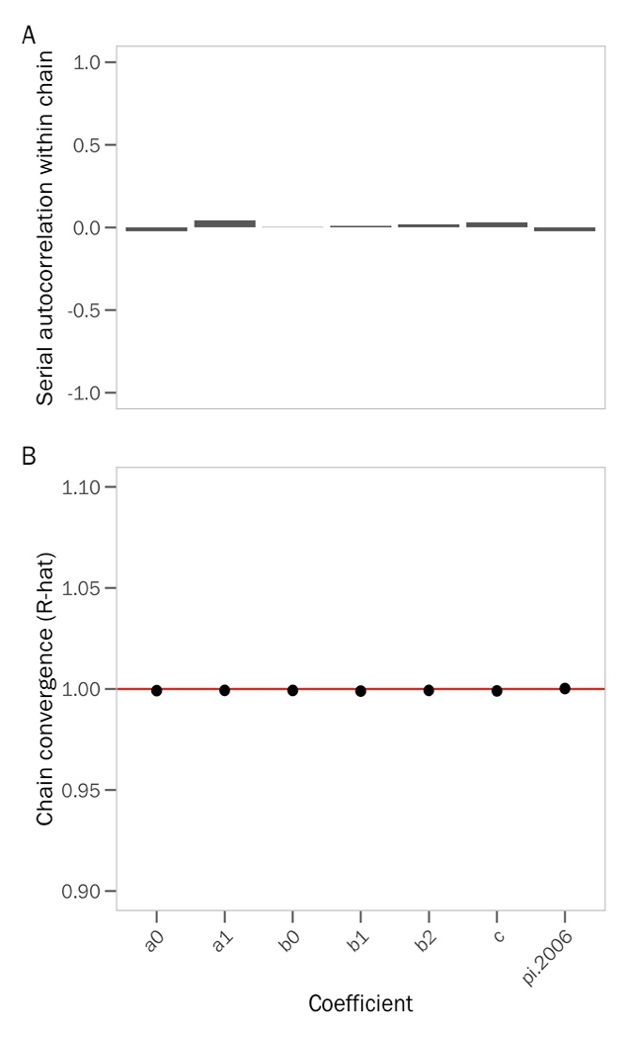

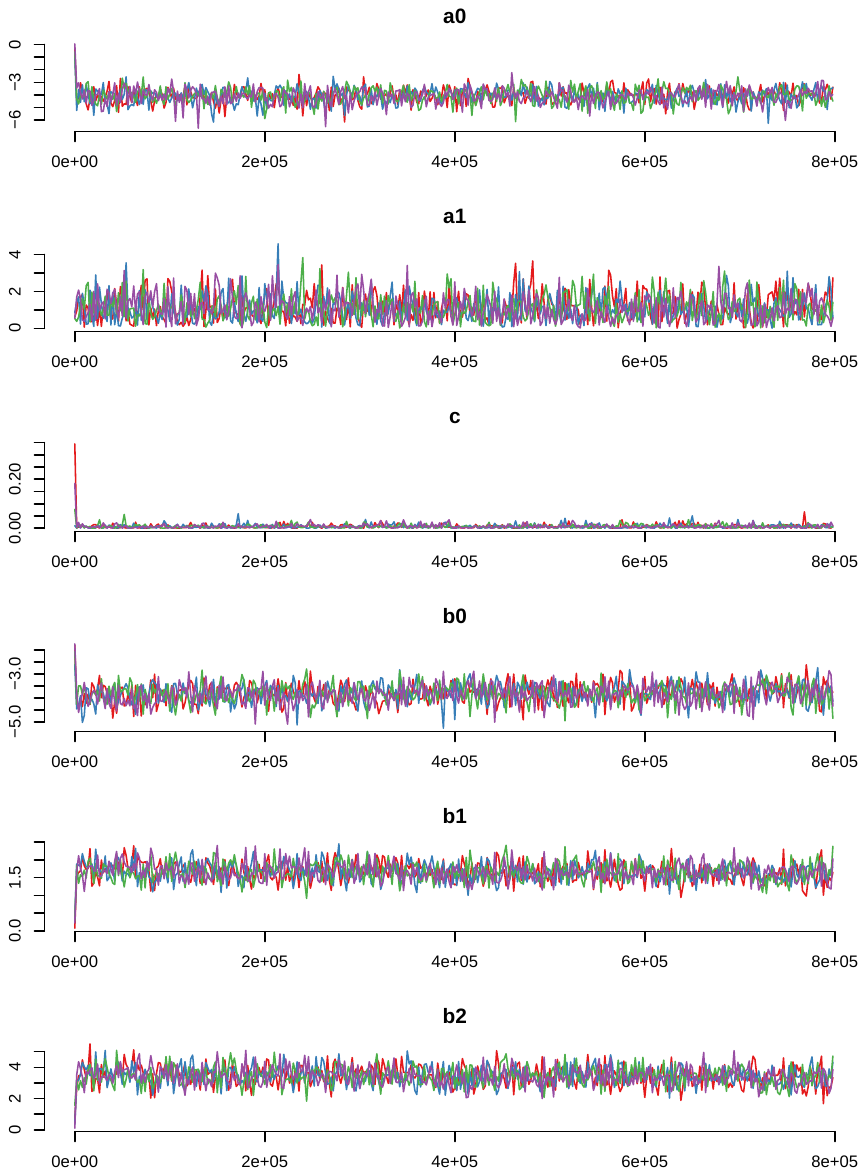

**Supplementary Text B**

*Considering the effect of individual heterogeneity in post-marking processes on mark-recapture models of survival*

Time-since-marking models control for losses to mortality or permanent emigration that occur during the interval after the first capture occasion (Pradel *et al.* 1997). To code a time-since-marking variable into the design matrix of the mark-recapture model, a time-varying covariate needs to be added that distinguishes the first occasion from all subsequent occasions. Unfortunately, time-varying covariates are unable to be implemented in the latest version of BaSTA, so we evaluated a time-since-marking Cormack–Jolly–Seber (CJS) model with the R package “RMark” (Laake 2013) and demographic parameters were estimated via maximum likelihood implemented in program MARK (White & Burnham 1999). We limited this analysis to unknown-aged individuals (i.e., only included individuals that were first captured as adults) because the first-year survival probability for individuals first marked as juveniles is 100% in our dataset because our dataset is limited to adults that have breeding information.

First, we evaluated the best model capturing variation in recapture probability while keeping apparent survival constant. In addition to the time-since-marking model, we tested an intercept only model, and a factorial time (i.e., year) model. This revealed that the factorial time model was far superior to the other models tested based on AICc statistics (Table S-B1). Furthermore, a comparison between the time-since-marking model and the intercept-only model shows that the DeltaAICc scores differ by <2 units – providing additional support that the time-since-marking model does not improve model fit. We interpret this as encouraging evidence that our inference about recapture probability is not hindered by individual heterogeneity in post-marking processes.

Second, we evaluated the if there was evidence of transience in our mark-recapture dataset that could bias our ability to appropriately estimate survival probability. In RMark we tested a time-since-marking survival model that incorporated annual variation in encounter probability structure shown above (i.e., Phi(~tsm)p(~time), Table S-B1). The results of this analysis revealed no difference in the survival probability of unknown-aged adults transitioning from their first or between subsequent capture occasions (Figure S-B1). Importantly, the estimated survival probability of the first occasion is actually larger than that of subsequent occasions (confidence intervals aside), indicating that there is little evidence that transient adults are biasing our mark-recapture analyses.

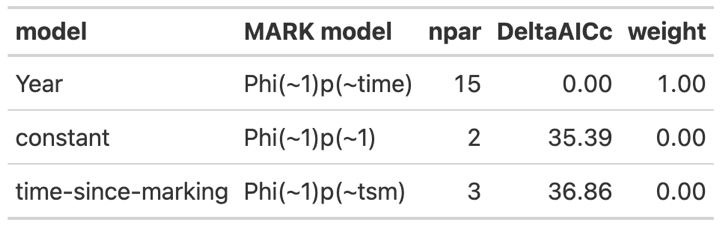

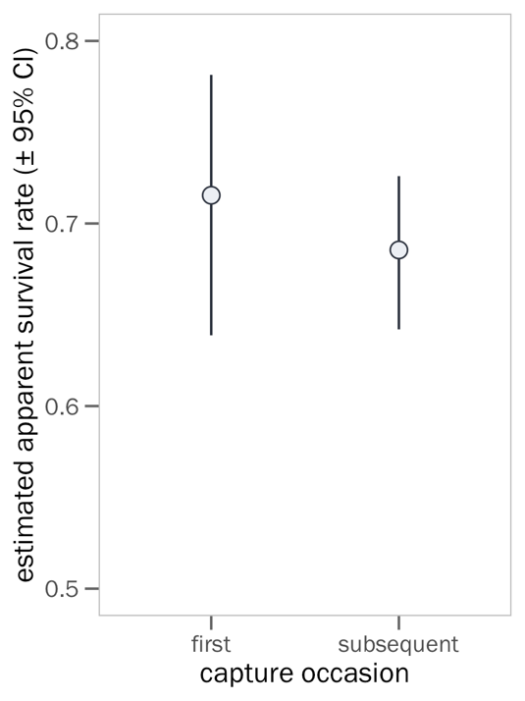
**Table S-B1.** AIC comparisons for three CJS models evaluating sources of variation in encounter probability. A time-since-marking model did not improve model fit over an intercept-only model, suggesting that our inference about recapture probability based on the BaSTA analyses (i.e., Supplementary Text A) are not hindered by individual heterogeneity in post-marking processes**.**

**Figure S-B1.** Results of a time-since-marking survival model that incorporated annual variation in encounter probability revealed no evidence that transient adults are biasing our mark-recapture-based inference.

**Supplementary Text C**

In theory, the selective benefits of larger eggs is that the subsequent hatchlings will be larger and have higher survival owing to more intrinsic reserves provided by the mother (Blomqvist *et al.*, 1997). To link egg size variation to potential fitness consequences of subsequent offspring we evaluated the predicted positive relationship between egg volume and chick weight using the egg dataset described above but reduced observations to the nest level and filtered to only include nests that had chicks measured within one day of hatching, resulting in 456 nests from 276 females. As it was unclear which chick came from which egg, each datum represented the nest-level average of chick weights and egg volumes. We included random intercepts for mother identity and year, and assumed a Gaussian error distribution of egg volume.

**Supplementary Text D**

*Considering the effect of body size on female survival*

To evaluate if individual quality predicts survival, we included tarsus length as a covariate in the RMark mark-recapture model (as described in Supplementary Text B above, individual covariates are not able to be incorporated into the latest version of BaSTA). Our analysis showed that there was no effect of tarsus length on survival probability (Figure S-D1). An intercept-only model had > 2 DeltaAICc units than the tarsus model (Table S-D1) and the graphical relationship between tarsus and survival unconvincing (Figure S-D1).

**Table S-D1.** AIC comparisons of two CJS models evaluating the effect of female tarsus length on apparent survival, while acknowledging annual variation in encounter probability.

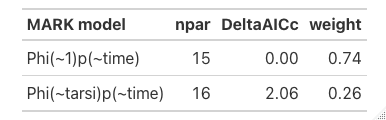

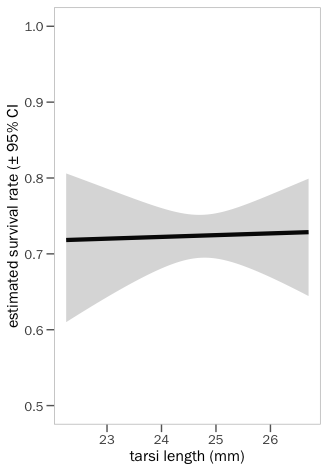
**Figure S-D1.** Model predictions for a relationship between apparent survival and female tarsus length reveal no relationship, while controlling for imperfect detection and annual variation in encounter probability.

**SUPPLEMENTARY FILES**

**Supplementary File 1.** Commented RMarkdown vignette detailing the analytical steps needed to reproduce all our models and results. Click here to view on your internet browser:

<https://raw.githack.com/leberhartphillips/snowy_plover_eggs/main/Rmd/Supplementary_File_1/Supplementary_File_1_.html>
